# Starvation of the bacterium *Vibrio atlanticus* induces simultaneous attacks on the dinoflagellate *Alexandrium pacificum*

**DOI:** 10.1101/2024.12.18.629110

**Authors:** Jean Luc Rolland, Estelle Masseret, Mohamed Laabir, Guillaume Tetreau, Benjamin Gourbal, Anne Thebault, Eric Abadie, Alice Rodrigues-Stien, Carole Veckerlé, Elodie Servanne-Meunier, Delphine Destoumieux-Garzón, Arnaud Lagorce, Raphaël Lami

**Author notes:** These authors contributed equally (Arnaud Lagorce, Raphaël Lami). Current address (Jean Luc Rolland). Corresponding author: Jean Luc Rolland, **Email:**.

## Abstract

Phytoplankton serve as a source of nutrients for bacteria in the marine environment. The interactions between algae and bacteria are known to include mutualism, commensalism, competition or antagonism. This occurs in the microenvironment surrounding phytoplankton cells, the phycosphere, an interface rich in nutrients and organic molecules exuded by the cell. Here, based on *in situ* observations and on an *in vitro* interaction study, we report on a novel form of starvation-induced hunting that the cells of selected Vibrio species exert on dinoflagellates. The results showed that *Vibrio atlanticus* was capable of attacking and killing the dinoflagellate *Alexandrium pacificum* ACT03. Briefly, the observed mechanism of algal-killing consists of first, the ‘immobilization stage’ involving the secretion of algicidal metabolites that disrupt the flagella of the algae. In the ‘attack stage’, Vibrios simultaneously surround algal cells at high density for a brief period without invading them. Finally, the ‘killing stage’ in which the lysis and consumption of the dinoflagellates occur. By using a combination of biochemical, proteomic, molecular and fluorescence microscopy approaches, we showed that this relationship is not related to the decomposition of algal organic matter, Vibrio quorum sensing pathways, toxicity of the algae or pathogenicity of the bacterium but is conditioned by nutrient stress, iron availability and linked to the iron-vibrioferrin transport system of *Vibrio atlanticus*. This is the first evidence of a new mechanism that could be involved in regulating Alexandrium spp. blooms and giving Vibrio a competitive advantage in obtaining nutrients from the environment. The interaction model we propose here suggests that Vibrio could play a role in regulating the proliferation of Alexandrium spp., giving it a competitive advantage in obtaining nutrients from the environment.

## INTRODUCTION

Harmful algal blooms (HABs) have experienced an increase in their occurrence, intensity, and geographical distribution on a global scale, resulting in adverse environmental, health, and socioeconomic impacts (Marampouti et al., 2021). HABs have a considerable impact on human health as a result of direct exposure to volatile toxins or by toxic seafood consumption (Burkholder et al., 2018). From an ecological point of view, the expansion of HABs can result in the erosion of biodiversity, because they cause massive mortality of marine species and they are generally monospecific in nature (Chai et al., 2020). In coastal areas, understanding the biological interactions that control toxic algal blooms is therefore a major ecological challenge. Among HAB-causing organisms, a number of *Alexandrium* species have been placed on the list of invasive Mediterranean species. Among them, *Alexandrium pacificum* is a flagellated eukaryotic unicellular organism that together with *Alexandrium tamarense* and *Alexandrium fundyense* forms the "Alexandrium tamarense" complex responsible for paralytic shellfish poisoning worldwide (Hadjadji et al., 2020). Since 1998, *A. pacificum* (former *A. catenella*) was monitored by the French phytoplankton observation and monitoring network (Rephy) in the Thau lagoon (French Mediterranean) because it produces paralytic shellfish toxins (PSTs) resulting in paralytic shellfish poisoning (PSP) syndrome.

Laanaia (Laanaia et al., 2013) showed that in Thau lagoon, a water temperature around 20°C for several days and organic and inorganic nutrients in sufficient concentrations are parameters favoring the development of *A. pacificum*, whose massive blooms occur in autumn. Algal blooms are seasonal events resulting in a rapid increase in the concentration of a species of algae in an aquatic environment. Depending on the species of phytoplankton, tolerance to physicochemical parameters varies, which influences when blooms occur (Leblad et al., 2020). Interestingly, although the collapse of phytoplankton blooms has been previously attributed to viruses (Pal et al., 2020), some ecological studies have suggested an important role of algicidal bacteria (Su et al., 2007; Wang et al., 2010). Among them several are belonging to the Vibrio genus (Li et al., 2014; Wang et al., 2020).

Vibrio (class γ-proteobacteria) are common microorganisms in marine systems worldwide (Baker-Austin et al., 2017; Mavian et al., 2020), where they are important components of the food chain, particularly in biodegradation, nutrient regeneration and biogeochemical cycles (Oberbeckmann et al., 2012). Vibrio is one of the most studied bacterial taxa due to their ubiquity in coastal marine systems and their capacity to cause infections in humans and animals, leading sometimes to epizootic or zoonotic epidemics (LeRoux et al., 2015; Mavian et al., 2020). Vibrio are extremely adaptable to their environment (Johnson, 2013). The main factors influencing their occurrence and distribution in water are temperature, salinity, nutrient availability (Wang et al., 2020), multiple strategies such as biofilm formation on biotic and abiotic surfaces (Espinoza-Vergara et al., 2020), or interactions with a multitude of other organisms such as eukaryotic predators (Drebes Dörr and Blokesch, 2020) or plankton (Lopez-Joven et al., 2018) are used by Vibrio in the environment. There is also evidence that global climate change has increased Vibrio-associated illnesses affecting humans and animals (Brumfield et al., 2021; Muhling et al., 2017). However, the drivers and dynamics of Vibrio survival and propagation in the marine environment are not yet fully understood.

A substantial number of research articles have highlighted the potential of γ-proteobacteria to exert algicidal activity against dinoflagellates, supporting the hypothesis that γ-proteobacteria such as Vibrio play a role in the control of algal blooms *in situ* (Coyne et al., 2022). However, the mechanisms behind Vibrio-driven algal lysis in the environment remain to be elucidated. Particularly, it is unclear how in the water column, algicidal compounds secreted by bacteria can concentrate around the algae to exert their lytic effect.

This study aims to describe observations made in the natural environment between Vibrio bacteria and Alexandrium algal blooms, and to determine *in vitro* the main factors involved in this relationship. Using a combination of biochemical, proteomic, molecular, and fluorescence microscopy approaches, we explored the role in algal toxicity, bacterial pathogenicity and the quorum sensing pathway on this relationship and showed the important role of nutrient stress and the iron uptake pathway in this unique Vibrio/Alexandrium interaction.

## METHODS

### Quantification of *Alexandrium* algae and Vibrio bacteria in the environment by qPCR

Seawater samples were collected in the Thau Lagoon (southern France, a shallow Mediterranean ecosystem open to the sea (Abadie et al., 1999), during spring and autumn 2015. Briefly, samples were collected from the subsurface (-50 cm) near an oyster table at a phytoplankton surveillance site (part of the REPHY network, N 43°26.058’ and E 003°39.878’). Once a week during spring and autumn 2015, during field sampling campaigns, 20 L of water was filtered on board through a 180 μm pore-size nylon membrane. At the laboratory, according to Lopez-Joven et al. (Lopez-Joven et al., 2018) seawater was fractionated into two size classes as follows: 2 L of the above filtrate was filtered through a 0.8 μm pore-size polycarbonate Whatman Nuclepore membrane to obtain organisms in the 0.8–180 μm range corresponding to plankton-associated Vibrio and living *Alexandrium* forms. Then, the filtrate from the 0.8 µm membrane was filtered again through a 0.2 μm pore-size polycarbonate Whatman Nuclepore membrane until the membrane was saturated. *Alexandrium* cells, ranging from 25 to 40 µm, belong to the microphytoplankton and are therefore retained in the 0.8–180 µm fraction. Any Vibrio cells potentially associated with or attached to *Alexandrium* cells will also be retained in this fraction. Vibrio cells are approximately 0.5-0.8 µm thick. The fraction between 0.2 and 0.8 µm therefore includes the free-living Vibrio. The bacterial population collected on 0.8-µm-pore-size filters was designated the particle-associated community, and the population on 0.2-µm-pore-size filters was designated as the free-living community. Membranes (in triplicate) were then conserved in 500 µL of 100% EtOH at -20°C. Environmental DNA (eDNA) was extracted from the MF Millipore membrane using the Macherey-Nagel NucleoSpin Tissue Kit and resuspended in 100 µL of water. The samples were then stored at -20°C after eDNA quantity and purity were assessed using a NanoDrop system (NanoDrop Technologies, Wilmington, DE, USA). PCR amplification reactions were done on a Roche LightCycler 480 Real-Time thermocycler (qPHD platform, University of Montpellier, France) using specific primer pairs (Table 3). Typically, the reactions contained 1 µL of template DNA (the DNA concentration for all samples varied from 1 to 40 μg mL^−1^), 0.5 µL of each primer (3.33 µM) and 4 µL of reaction mixture (SYBR Green Master Mix) in a total volume of 6 μL. The reaction parameters were as follows: 5 minutes at 95°C (initial denaturation) and 40 cycles of 10 s at 95°C (denaturation), 10 s at the corresponding hybridization temperature (Table 3) and 10 s at 72°C (elongation). Melting curve profiles were generated by increasing the temperature from 65°C to 95°C at 0.5°C s^-1^. Amplification products were analysed using LightCycler software (Roche Diagnostics). *Vibrio spp*. and *A. pacificum* and *A. tamarense* were quantified by constructing calibration curves based on DNA from the *V. atlanticus* LGP32 reference strain (former *V. tasmaniensis* LGP32) and from the *A. pacificum* reference strain (ACT03: *A. catenella* strain isolated from Thau in 2003) and the *A. tamarense* reference strain (ATT07: *A. tamarense* isolated from Thau in 2007) (not shown).

### Strains and growth conditions

Vibrio strains. Wild-type and isogenic mutants of *V. atlanticus* LGP32 (Table 1) were used in this study. Deletion-mutants included Δ*luxR*, Δ*luxS*, Δ*luxM* and Δ*pvuB* isogenic strains. The Δ*pvuB* mutant was constructed here by allelic exchange as described previously by Le Roux (Le Roux et al., 2007). We also used *V. atlanticus* LGP32 carrying the pSW3654T-GFP plasmid (Le Roux et al., 2007), hereafter referred to as *V. atlanticus* LGP32-GFP. Bacterial strains were grown at 22 ± 1°C in Zobell medium (0.38 µM iron (III)). When needed, 25 μg mL^-1^ chloramphenicol (Cm) was added to cultures of *V. atlanticus* LGP32-GFP (Le Roux et al., 2007).

**Table 1.** Ability of Vibrio strains to attack and to lyse *Alexandrium pacificum*. ND: not determined.

| **Vibrio species** | **Strains** | **Virulence for fish or invertebrates** | **References** | **Attack *A. pacificum*** | **Lyse *A. pacificum*** |
| --- | --- | --- | --- | --- | --- |
| *Vibrio atlanticus* LGP32 | WT | Yes | (Gay et al., 2004) | + | + |
| // | WT + pSW3654T-GFP | Yes | (Le Roux et al., 2007) | + | + |
| // | ΔLuxM | ND | Ifremer Institute, France | + | + |
| // | ΔLuxS | ND | // | + | + |
| // | ΔLuxR | ND | // | + | + |
| // | ΔPvuB | ND | This work | - | + |
| *Vibrio tasmaniensis* | J5-9 | Yes | (Lemire et al., 2015) | + | + |
| // | LMG20012<sup>T</sup> | No | (Thompson et al., 2003) | + | + |
| *Vibrio crassostreae* | J2.9 | Yes | (Lemire et al., 2015) | + | + |
| // | J2-8 | No | // | + | + |
| *Vibrio fischeri* | ES114 | ND | (Mandel et al., 2008) | + | + |
| *Vibrio harveyi* | ATCC14126 | Yes | (Liu et al., 1996) | + | + |
| *Vibrio aestuarianus* | janv-32 | Yes | (Labreuche et al., 2010) | + | + |

*Phytoplankton strains*. Non-axenic phytoplankton species (Table 2) were grown in batch culture in enriched natural sea water (ENSW, 6.55 µM iron (III)) with a salinity of 36 practical salinity units (PSU) at 22 ± 1°C under cool white fluorescent illumination (100 µmol photons m^-2^ s^-1^) and a 12 h:12 h light:dark cycle (Harrison et al., 1980). The algae were used for experiments in their exponential growth phase.

**Table 2.**
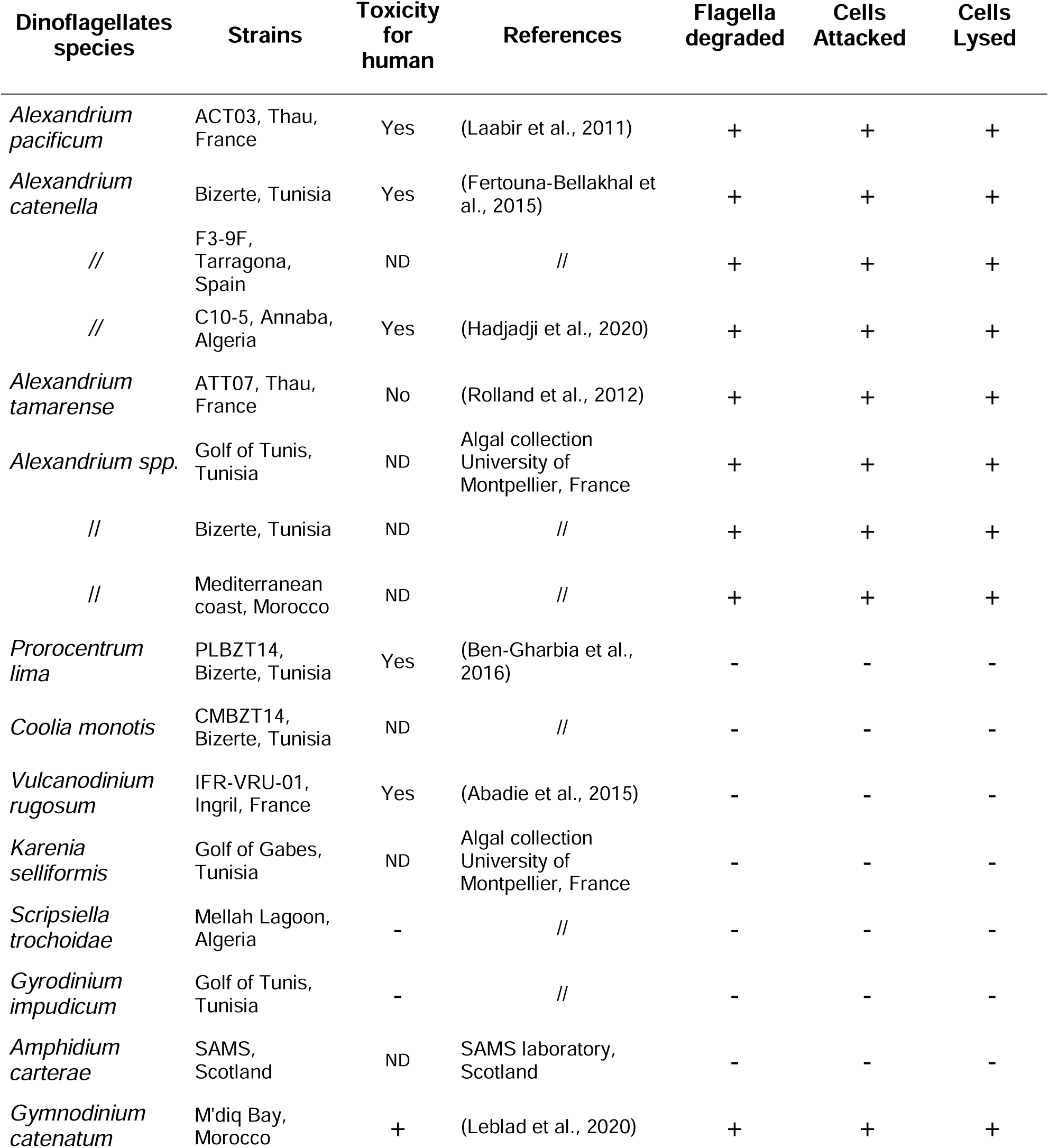
Ability of *V. atlanticus* LGP32 to degrade flagella, attack and lyse the targeted dinoflagellates *spp*. commonly found in the Mediterranean Sea. ND not determined.

| Dinoflagellates species | Strains | Toxicity for human | References | Flagella degraded | Cells Attacked | Cells Lysed |
| --- | --- | --- | --- | --- | --- | --- |
| *Alexandrium pacificum* | ACT03, Thau, France | Yes | (Laabir et al., 2011) | + | + | + |
| *Alexandrium catenella* | Bizerte, Tunisia | Yes | (Fertouna-Bellakhal et al., 2015) | + | + | + |
| // | F3-9F, Tarragona, Spain | ND | // | + | + | + |
| // | C10-5, Annaba, Algeria | Yes | (Hadjadji et al., 2020) | + | + | + |
| *Alexandrium tamarense* | ATT07, Thau, France | No | (Rolland et al., 2012) | + | + | + |
| *Alexandrium spp.* | Golf of Tunis, Tunisia | ND | Algal collection University of Montpellier, France | + | + | + |
| // | Bizerte, Tunisia | ND | // | + | + | + |
| // | Mediterranean coast, Morocco | ND | // | + | + | + |
| *Prorocentrum lima* | PLBZT14, Bizerte, Tunisia | Yes | (Ben-Gharbia et al., 2016) | - | - | - |
| *Coolia monotis* | CMBZT14, Bizerte, Tunisia | ND | // | - | - | - |
| *Vulcanodinium rugosum* | IFR-VRU-01, Ingril, France | Yes | (Abadie et al., 2015) | - | - | - |
| *Karenia selliformis* | Golf of Gabes, Tunisia | ND | Algal collection University of Montpellier, France | - | - | - |
| *Scropsiella trochoidea* | Mellah Lagoon, Algeria | - | // | - | - | - |
| *Gyrodinium impudicum* | Golf of Tunis, Tunisia | - | // | - | - | - |
| *Amphidium carterae* | SAMS, Scotland | ND | SAMS laboratory, Scotland | - | - | - |
| *Gymnodinium catenatum* | M'diq Bay, Morocco | + | (Leblad et al., 2020) | + | + | + |

### Co-culture assay

For each tested phytoplankton species (Table 2), 2 x 10^4^ cells harvested in their exponential growth phase (doubling time between 5 and 7 days) were placed in 20 mL of ENSW medium in a 50 mL suspension culture flask (Cellstar® PS, Greiner bio-one). After incubation for 24 h at 22 ± 1°C under cool white fluorescent illumination, 40 µL of Vibrio strains (Table 1) grown for 12, 36, 60 and 156 h in Zobell medium or in Zobell medium supplemented with FeCl_3_ (6 µM iron (III)) or with boron (H_3_BO_3;_ 0.47 mM) or the corresponding culture supernatant were added to the phytoplankton cells (Table 2). After incubation at 22 ± 1°C for 0, 15, 30, 45 and 60 min under cool white fluorescent lights, living, non-swimming, attacked and lysed phytoplankton cells were counted in a sedimentation chamber under an inverted microscope. The number of lysed cells corresponded to phytoplankton cells showing disrupted membranes. Non-swimming algae were not counted as lysed cells. For Vibrio analysis, 100 µL of a 1:10 serial dilution mixture in ENSW (from 10^-2^ to 10^-10^) was plated on Vibrio Selective TCBS (thiosulfate-citrate-bile salts-sucrose) agar (in triplicate). After incubation for 24 h at 22 ± 1°C, the number of living Vibrio cells was determined by counting colony-forming units (CFUs). The data come from three independent experiments using independent phytoplankton cultures and independent bacterial cultures.

### Microscope observations

The dynamic of the interaction between *A. pacificum* ACT03 isolated from the French Thau Lagoon, south of France (Laabir et al., 2011) and *V. atlanticus* LGP32, which is an oyster pathogen isolated from the French Atlantic coast (Gay et al., 2004) and present in the Thau Lagoon (Lopez-Joven et al., 2018) was surveyed. As the *A. pacificum* ACT03 strain (Table 2) used in the study is not axenic, there is potential for bacteria other than *V. atlanticus* LGP32 to be present in the experiments. To elucidate the interaction without thoroughly accounting for the non-axenic cells, interaction was observed under a Zeiss Axio upright fluorescence microscope equipped with an AxioCamMRm 2 digital microscope camera using *V. atlanticus* LGP32 tagged with green fluorescent protein (GFP). Lasers were used at excitation wavelength (λex) 488 nm for GFP (emission wavelengths (λem): 505–530 nm) and λex 532 nm for plankton chloroplasts (λem 560–630 nm). Images were taken sequentially to avoid cross-contamination between fluorochromes. Sequences of images were merged during the Vibrio-*Alexandrium* interaction using ZEN 2012 (blue edition) software. Interaction events between Vibrio strains and phytoplankton strains were also observed under a Leica TCS SPE confocal laser scanning system connected to a Leica DM 2500 upright microscope camera (Montpellier RIO Imaging Platform, University of Montpellier, France).

### Comparative proteomic analysis

#### Vibrio sampling and protein extraction

*V. atlanticus* LGP32 was grown for 60 h at 22°C in artificial seawater (high nutrient stress, cond. 1) or 12 h at 22°C in Zobell media (no nutrient stress, cond. 2). After 10 min centrifugation at 8000 rpm, crude protein extracts of *V. atlanticus* LGP32 in each culture condition (triplicates) were obtained by sonication on ice at 20% amplitude for 20 s in 200 µL of ice-cold denaturing buffer (7 M urea, 2 M thiourea, 4% CHAPS in 30 mM Tris-HCl, pH 8.5) and clarified by centrifugation at 2000 x *g*, 15 min, 4°C. The protein concentration of the supernatant was estimated using the 2D Quant Kit (Cytiva™, MERCK) and samples were stored at -80°C until use.

#### Two-dimensional gel electrophoresis (2D gel)

Protein extracts were individually analysed on 2D gel electrophoresis (6 gels per condition each corresponding to different biological replicates). To do so, 100 µg of proteins from each extract was added to rehydration buffer (7 M urea, 2 M thiourea, 4% CHAPS, 65 mM DTT) for a total volume of 350 µL. They were then individually loaded onto 17 cm isoelectric focusing strips (Bio-Rad) with a stabilized non-linear pH ranging from 3 to 10. Due to the high complexity of the protein profile in the acidic part (left) of the gel pH 3–10 (Fig. S3), we conducted additional ‘close-up’ analyses in gels using 17 cm isoelectric focusing strips (Bio-Rad) with a narrower, stabilized pH gradient ranging from 4 to 7. Strips were rehydrated passively for 5 h at 22°C, followed by active rehydration for 14 h under a 50 V current at 22°C (to help large proteins enter the strips). Thereafter, isoelectrofocusing was carried out using the following programme: 50 V for 1 h, 250 V for 1 h, 8000 V for 1 h and a final step at 8000 V for a total of 140 000 V.h with a slow ramping voltage (quadratic increasing voltage) at each step. Focused proteins were reduced by incubating the strip twice in equilibration buffer (1.5 M Tris, 6 M urea, 2% SDS, 30% glycerol; bromophenol blue, pH 8.8) containing DTT (130 mM) at 55°C. Then, they were alkylated by incubation with equilibration buffer containing iodoacetamide (135 mM) on a rocking agitator (400 rpm) at room temperature protected from light. Proteins were also separated according to their molecular weight (second dimension) on 12% acrylamide/0.32% piperazine diacrylamide gels run at 25 mA per gel for 30 min followed by 75 mA per gel for 8 h using a Protean II XL system (Bio-Rad). Gels were stained using an MS-compatible silver staining protocol and scanned using a ChemiDoc MP Imaging System (Bio-Rad) associated with Image Lab software version 4.0.1 (Bio-Rad).

#### Comparative bioinformatics analysis of 2D gels

Twelve gels (six per condition) were selected for comparative analysis on PD-Quest v. 7.4.0 (Bio-Rad) to identify changes in protein abundance between the proteomic profiles of *V. atlanticus* LGP32 cultured in contrasting nutrient conditions (ENSW/Zobell). Spots whose mean intensity across six replicates per strain was two times higher or lower than those from the other strain, with a *P* <0.01 (Mann-Whitney U-test), were considered significantly different in terms of abundance between the two conditions (quantitative difference). Differentially represented spots were then excised from the gels, destained, trypsin-digested and the obtained peptides were identified by tandem mass spectrometry (MS-MS) using the PISSARO platform facility (University of Rouen, France). To identify protein(s) present in each spot, the obtained peptides were compared with *V. atlanticus* LGP32 reference genome (https://vibrio.biocyc.org/). The genes whose peptides matched strongly were retrieved and used for a BLASTx query against non-redundant databases to determine the protein identity of the best match. A gene was considered as strongly matched when at least two peptides matched the sequence with a coverage of > 6%. Their theoretical isoelectric point (pI) and molecular weights were also calculated using the Expasy server (https://www.expasy.org/) to compare them with the location of the spot on the gel. Altogether, these complementary analyses made it possible to characterize the protein identity of each spot with confidence.

### Gene expression analysis

#### Vibrio sampling and RNA extraction

*V. atlanticus* LGP32 was grown in Zobell media for 36 h and 60 h at 22°C (decline phase of growth, nutrient stress) or for 12 h at 22°C (exponential growth phase, no nutrient stress). Total RNA was isolated from *V. atlanticus* LGP32 using the standard TRIzol method (Invitrogen Life Technologies SAS, Saint-Aubin, France) and then treated with DNase (Invitrogen) to eliminate genomic DNA contamination. After sodium acetate precipitation, the quantity and quality of the total RNA were determined using a NanoDrop spectrophotometer and agarose gel electrophoresis. Following heat denaturing (70°C for 5 minutes), reverse transcription was performed using 1 μg of RNA prepared with 50 ng μL^-1^ oligo-(dT) 12–18 mer in a 20-μL reaction containing 1 mM dNTPs, 1 unit μL^-1^ RNAseOUT and 200 units μL^-1^ Moloney murine leukaemia virus reverse transcriptase (M-MLV RT) in reverse transcriptase buffer, according to the manufacturer’s instructions (Invitrogen Life Technologies SAS, Saint-Aubin, France).

#### PCR amplification

Amplification reactions were analysed using a Roche LightCycler 480 Real-Time thermocycler (Bio-Environnement platform, University of Perpignan, France). In this study, several PCR primer pairs were designed using Primer3 software (optimal primer size: 20 bases; Tm: 60°C; primer GC%: 50; 2GC clamp and product size range: 150–200 bp) and calibrated with *V. atlanticus* LGP32 genomic DNA (Table 1). To determine the qPCR efficiency of each primer pair used, standard curves were generated using seven serial dilutions of genomic DNA (10^-2^, 10^-3^, 10^-4^, 10^-5^, 10^-7^ and 10^-8^) (not shown); the qPCR efficiencies of the tested genes varied between 1.85 and 2.08 (Table 3). For gene expression, reverse transcription was performed with 1 µg of total RNA using random hexamers and SuperScript IV reverse transcriptase (Invitrogen). The total qPCR reaction volume was 10 μL and consisted of 5 μL of cDNA (diluted 1:5), 2.5 μL of SensiFAST SYBR No-ROX Mix (Bioline) and 100 nM or 300 nM PCR primer pair (Table 3). The reaction parameters were as follows: 2 min at 95°C (initial denaturation) and 40 cycles of 5 s at 95°C (denaturation), 10 s at 59°C (annealing) and 20 s at 72°C (elongation). The specificity of each PCR was checked by measuring fluorescent signals during melting curve analysis (PCR product heated from 65°C to 95°C continuously and slowly at 0.1°C s^−1^). Relative expression was calculated by normalization to the expression of two constitutively expressed housekeeping genes, namely, 6PKF (VS_2913) and CcmC (VS_0852), using the delta-delta threshold cycle (ΔΔCt) method (Pfaffl, 2001).

**Table 3.**
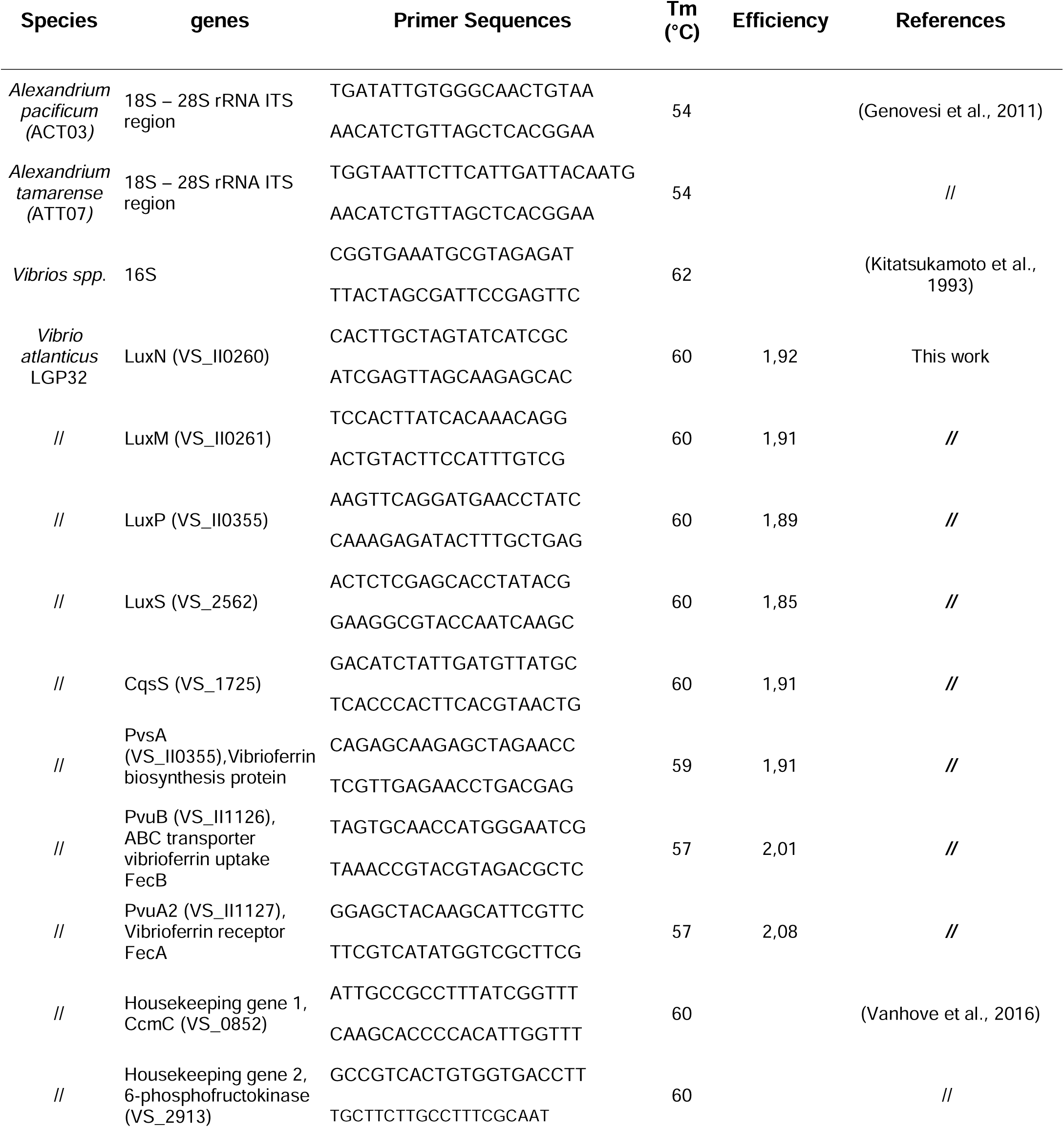
Oligonucleotide sequences of primers used for RNA expression analysis.

### Detection of quorum-sensing signalling molecules

#### Vibrio culture

To detect the QS molecules (AI-2, AI-1 and CAI-1) *V. atlanticus* LGP32 was grown in Zobell media for 12 h (exponential growth phase, control) and 60 h (decline phase of growth, nutrient stress).

#### AI-2 analysis

Bioluminescence assay using the QS bioluminescent of *Vibrio campbellii* MM32 (*luxN*::Cm, *luxS*::Tn5Kan) was used to detect AI-2 molecules in culture supernatants. Briefly, *Vibrio* cultures were centrifuged at 17,000 x *g* for 10 min, and the resulting supernatants were filtered on 0.22 μm. Then 20 μL of the filtrates were mixed with 180 μL of *V. campbellii* MM32 diluted 1:5000 then incubated at 30°C and 100 rpm. Luminescence and cell density (OD620) were collected in triplicate and analysed according to Tourneroche et al.(Tourneroche et al., 2019).

#### AI-1 and CAI-1 extraction and LC-MS analysis

Chemical analyses were conducted with a Q Exactive Focus Orbitrap System coupled to an Ultimate 3000™ ultrahigh-performance liquid chromatography (UHPLC) system (Thermo Fisher Scientific) according to Rodrigues et al. (Rodrigues et al., 2022). Briefly, ethyl acetate (2 mL) was added into each culture (2 mL). This mixture was shaken overnight at room temperature (150 rpm). The two phases were then separated and the aqueous phase was extracted once again. The two obtained organic phases were pooled and the solvent was evaporated under vacuum. The crude extracts were dissolved in 500 µL LC-MS grade methanol for analysis. The experiments were performed using biological triplicates, each of which was analyzed in triplicate. Analyses of extracts and standards (3 μL injected) were performed in electrospray positive ionization mode in the 50–750 m/z range in centroid mode. The parameters were as follows: spray voltage: 3 kV; sheath flow rate: 75; aux gas pressure: 20; capillary temperature: 350°C; heater temperature: 430°C. The analysis was conducted in Full MS data-dependent MS2 mode (Discovery mode). Resolution was set to 70,000 in Full MS mode, and the AGC (automatic gain control) target was set to 1x10^6^. In MS2, resolution was 17,500, AGC target was set to 2x10^5^, isolation window was 0.4 m/z, and normalized collision energy was stepped to 15, 30 and 40 eV. The UHPLC column was a Phenomenex Luna Omega Polar C18 1.6 μm, 150 x 2.1 mm. The column temperature was set to 42°C, and the flow rate was 0.4 mL min^-1^. The solvent system was a mixture of water (A) with increasing proportions of acetonitrile (B), with both solvents modified with 0.1% formic acid. The gradient was as follows: 1% B 3 min before injection, then from 1 to 15 min, a gradient increase of B up to 100% (curve 5), followed by 100% B for 5 min. The flow was injected into the mass spectrometer starting immediately after injection. All data were acquired and processed using FreeStyle 1.5 software (Thermo Fisher Scientific).

#### Chemicals and solvents

N-acyl-homoserine lactones (AHL) were obtained from Cayman Chemical (Ann Arbor, MI, USA). Stock solutions were obtained by dissolving standards in methanol or dichloromethane (C18-AHL) at a concentration of 1 mg mL^-1^ and stored at -80°C. Standard solutions for UHPLC- high-resolution tandem mass spectrometry (HRMS) analyses were prepared by diluting each individual standard solution with methanol in order to obtain a concentration range from 2000 to 20 ng mL^-1^. LC-MS grade methanol, acetonitrile and formic acid were purchased from Biosolve (Biosolve Chimie, Dieuze, France), analytical-grade ethyl acetate was obtained from Sigma-Aldrich. Pure water was obtained from Elga Purelab Flex System (Veolia LabWater STI, Antony, France).

### Nature of lytic compounds secreted by *V. atlanticus* LGP32

To determine the temperature-sensitivity of the lytic compounds secreted, *V. atlanticus* LGP32 grown for 60 h in Zobell media at 22 °C was filtered through a 10 kDa membrane (Amicon^®^ Ultra-4 filter unit). The eluate containing molecules with MW below 10 kDa was then incubated in a water bath at 100°C for 30 min. Boiled filtrates (0.1 mL) were subsequently used to inoculate *A. pacificum* (ACT03 strain) cultures, then lytic activity was observed under the Leica TCS SPE confocal laser scanning system. Zobell media with the same treatment was used as control.

### Statistical analysis

#### Environmental data

Statistical analyses were performed using R 3.6.3 software (R Core Team, 2024) using Rstudio. The relationship between *Alexandrium* and Vibrio was explored separately in spring and autumn. We used a generalized linear model specifying a Gaussian family. For spring, the dataset for salinity and temperature was complete (14 periods of observation). An influential period was detected and removed from the dataset because no dead *Alexandrium* cells were observed. The effects of explanatory variables such as log10 (Vibrio+1), salinity and temperature were centred, reduced and tested as fixed effects with a linear relationship. Model selection was performed using the Akaike information criterion corrected for small sample size (AIC_c_). Models were considered different whenever the difference between their AIC_c_ value and the lowest AIC_c_ value (ΔAIC_c_) was lower than 2 (Burnham and Anderson, 2002). *Alexandrium* distribution and model residuals were checked for normal distribution assumptions (QQ plot and Shapiro-Wilk test). For autumn, the dataset was complete for 10 periods of observation. Salinity and temperature were missing for three periods. We explored the relationship between *Alexandrium* and Vibrio alone using the method detailed above for spring.

#### In vitro data

Statistical analyses were performed using one-way ANOVA (analysed by pair) followed by Tukey’s test (Statistica 10.0 software, StatSoft, Maison-Alfort, France). *P <0.05, **P <0.01, ***P <0.001.

## RESULTS

### Concomitant occurrence of *A. pacificum* ACT03*, A. tamarense* ATT07 and free-living *Vibrio spp.* in the Thau lagoon

In the spring and autumn of 2015, in the Thau Lagoon (Fig. 1A), we detected *Alexandrium* algae (*A. pacificum* ACT03 *and A. tamarense* ATT07, both alive and in degraded cell forms) and free-living Vibrio, but no plankton-associated Vibrio were observed (Fig. 1B, Data S1). Using model selection based on AICc, we found no significant relationship between *Alexandrium* (*A. pacificum* ACT03*, A. tamarense*) and *Vibrio spp.* abundances in autumn. This result is consistent with Vibrio’s difficulty in growing at temperatures below 20°C, as well as with the many environmental factors that can influence the dynamics of algae proliferation (Laanaia et al., 2013). Interestingly, in spring 2015, the mean densities of all *Alexandrium* cells (degraded and alive) and of free-living Vibrio were positively correlated. The lowest AICc was obtained with the model explaining degraded form of *Alexandrium* density based on the free Vibrio density (Fig. 1C). Given that, this model is not so different from the model with only the intercept, but better than any other linear combination with other potentially interfering drivers, such as temperature and salinity (Fig. 1D), we searched for evidence of a relationship between Vibrio and *Alexandrium* by studying their interaction *in vitro*.

**Figure 1.**
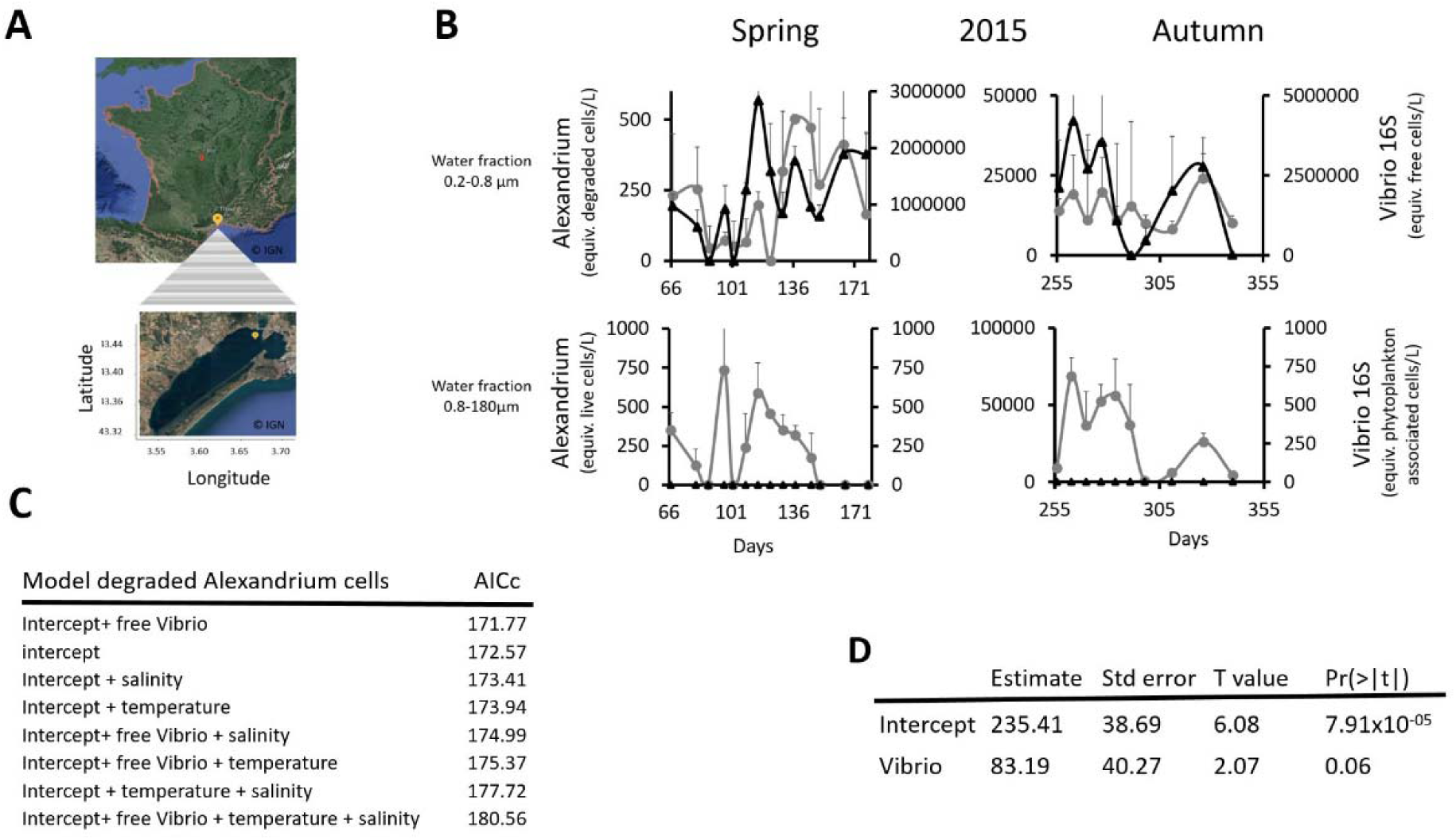
Dynamics of *Alexandrium* and Vibrio in the environment. **(A)** Location of the monitoring station in the Thau Lagoon (southern France). **(B)** Mean abundance (DNA equiv.) of *Vibrio spp*. (16S) and Alexandrium *spp*. *(A. pacificum* ACT03 *+ A. tamarense* ATT07). Vibrio cells (black line with diamond dot) and degraded *Alexandrium* cells (grey line with round dot) were detected in the 0.2–0.8 μm fraction (free Vibrio fraction) in spring and autumn 2015. Living *Alexandrium* cells (grey line with round dot) but no plankton-associated *Vibrio spp*. (black line with diamond dot) were evidence in the 0.8–180 μm in spring and autumn. **(C)** Results of the Akaike Information Criterion (AICc) test conducted to select a model for explaining the mean value of dead Alexandrium (degraded cells) in spring. **(D)** Wald test of the AICc model explaining the mean value of dead Alexandrium in spring by free Vibrio.

### V. atlanticus LGP32 feeds on Alexandrium pacificum ACT03

To investigate whether *Alexandrium* interacts with Vibrio, we incubated in Enriched Natural SeaWater (ENSW) *A. pacificum* ACT03 (2.0 x 10^4^ cells) with *V. atlanticus* LGP32 previously grown for 12 hours in Zobell media (initial concentration of 8.8 x 10^7^ cells mL^-1^). In interaction with *V. atlanticus* LGP32, *A. pacificum* ACT03 cell abundance decreased significantly from 2.1 x 10^4^ cells mL^-1^ after 1h of exposure to 1.1 x 10^4^ cells mL^-1^ after 48 h of exposure (Fig. 2A), while the *V. atlanticus* LGP32 concentration grew significantly after 26 h of interaction, reaching a maximum peak density of 7.6 x 10^7^ CFU mL^-1^ at 34 h (Fig. 2B). In the control experiment where *A. pacificum* ACT03 was cultured alone in ENSW, the algal concentration remained stable over time (Fig. 2A) and no bacteria were on the corresponding TCBS plates (Vibrio selective medium). In the control where *V. atlanticus* LGP32 was grown alone in ENSW, the bacterial concentration decreased from 7.0 × 10L CFU/mL after 1 hour of incubation to 1.1 × 10L CFU/mL after 48 hours of incubation (Fig. 2B). These results show that the interaction between *V. atlanticus* LGP32 and *A. pacificum* ACT03 leads to a decline in the algal population and promotes the growth of *V. atlanticus* LGP32. This suggests that *V. atlanticus* LGP32 is able to feed on *A. pacificum* ACT03.

**Figure 2.**
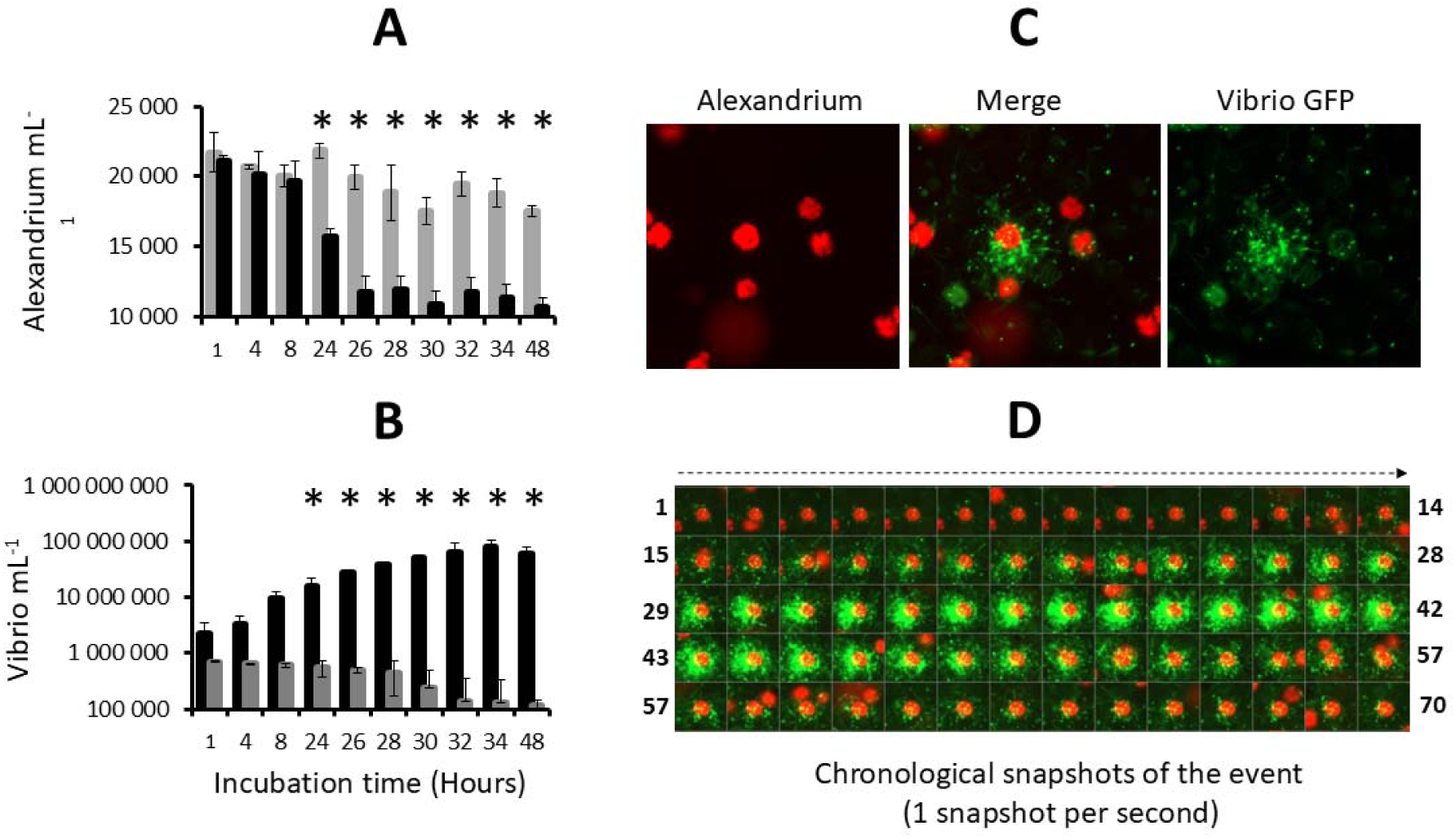
Incubation of *V. atlanticus* LGP32 and *Alexandrium pacificum* ACT03 in enriched natural seawater (ENSW). **(A)** *A. pacificum* ACT03 cultured alone (grey bar) and incubated with *V. atlanticus* LGP32 (black bar) in ENSW. **(B)** *V. atlanticus* LGP32 cultured alone (grey bar) and incubated with *A. pacificum* ACT03 (black bar) in ENSW. **(C)** Snapshot of the interaction between *V. atlanticus* LGP32-GFP cells (60-hour culture) and one cell of *A. pacificum* ACT03 taken at 8h00 of co-culture. **(D)** Chronological snapshots of the interaction (70 pictures, one per second). *V. atlanticus* LGP32 (small green cells) and *A. pacificum* ACT03 cell (large red cell). All experiments were done in triplicate. Asterisks indicate significant differences in a multiple comparison test (One-way ANOVA with post hoc Tukey test), *P ≤ 0.05.

### *V. atlanticus* LGP32 performs attacks on *A. pacificum* ACT03

Epifluorescence microscopy observation of GFP-labelled *V. atlanticus* LGP32 (previously grown in Zobell medium) in interaction showed that *V. atlanticus* LGP32 cells are capable of simultaneously attacking *A. pacificum* ACT03 cells (Fig. 2C and Video 1). The attacks were extremely rapid, with empty thecae (algal envelopes) observed in the medium after less than 60 s (Fig. 2D and Video 2). During the attack, *V. atlanticus* LGP32 did not invade the algal cell but remained clustered on the cell surface (Fig. 2C).

### Attack of *A. pacificum* ACT03 is activated by *V. atlanticus* LGP32 starvation

In 2002, Martin hypothesized that nutritional stress induces bacteria to lyse algae (Martin, 2002). To test this hypothesis, we monitored *V. atlanticus* LGP32 behaviour in response to starvation (Fig. 3). We observed that *V. atlanticus* LGP32 in exponential growth phase (12 h of culture in Zobell medium, 8.8 x 10^7^ cells mL^-1^) did not interact with *A. pacificum* ACT03 cells for the first hour of contact (Fig. 3A, B, C). In contrast, *V. atlanticus* LGP32 in the decline phase (36 h of culture in Zobell medium, 1.3 x 10^7^ cells mL^-1^) induced a significant decrease in the number of motile algae cells by 8.9% after 15 min and by 43.3% after 60 min (Fig. 3A). This phenomenon corresponded to the degradation and/or disruption of algal flagella (Video 3). The flagella no longer functioned correctly, which caused irregular swimming of the algae (Video 3, left cell). This was followed by a complete cessation of swimming. When the flagellum detached from the algae (Video 3, right cell), the attack occurred. With starved *V. atlanticus* LGP32 (60 h of culture in Zobell medium, 0.6 x 10^7^ cells mL^-1^), algae immobilization was fast and significant (91.4% in 15 min, Fig. 3A), and algae were attacked individually being targeted by *V. atlanticus* LGP32 cells (Video 4). The percentage of cells attacked and killed peaked at 30% after 15–30 min of contact (Fig. 3B and 3B1) and then decreased. After 1 h, attacks had stopped with approximately 40% of the algal cells still alive (Fig. 3B1 and 3C). Although it remains unclear whether the attacks occur during a specific phase of growth, it is evident that the cells are already weakened before attack as they have all lost their flagella. An old-starved culture of *V. atlanticus* LGP32 (126 h of culture in Zobell medium, < 0.1 x 10^7^ cells mL^-1^) significantly immobilized *A. pacificum* ACT03 cells within a few minutes, with lysis occurring immediately (Fig. 3A and 3C), making it impossible to detect attacks by *V. atlanticus* LGP32 (Fig. 3B). The lysis phase corresponded to initial vesicle formation followed by the bursting of *A. pacificum* ACT03 cells (Fig. 3C and 3C1). Importantly, Vibrio densities decreased with culture age, ruling out the possibility that the stronger predation observed in older cultures was driven by higher bacterial densities.

**Figure 3.**
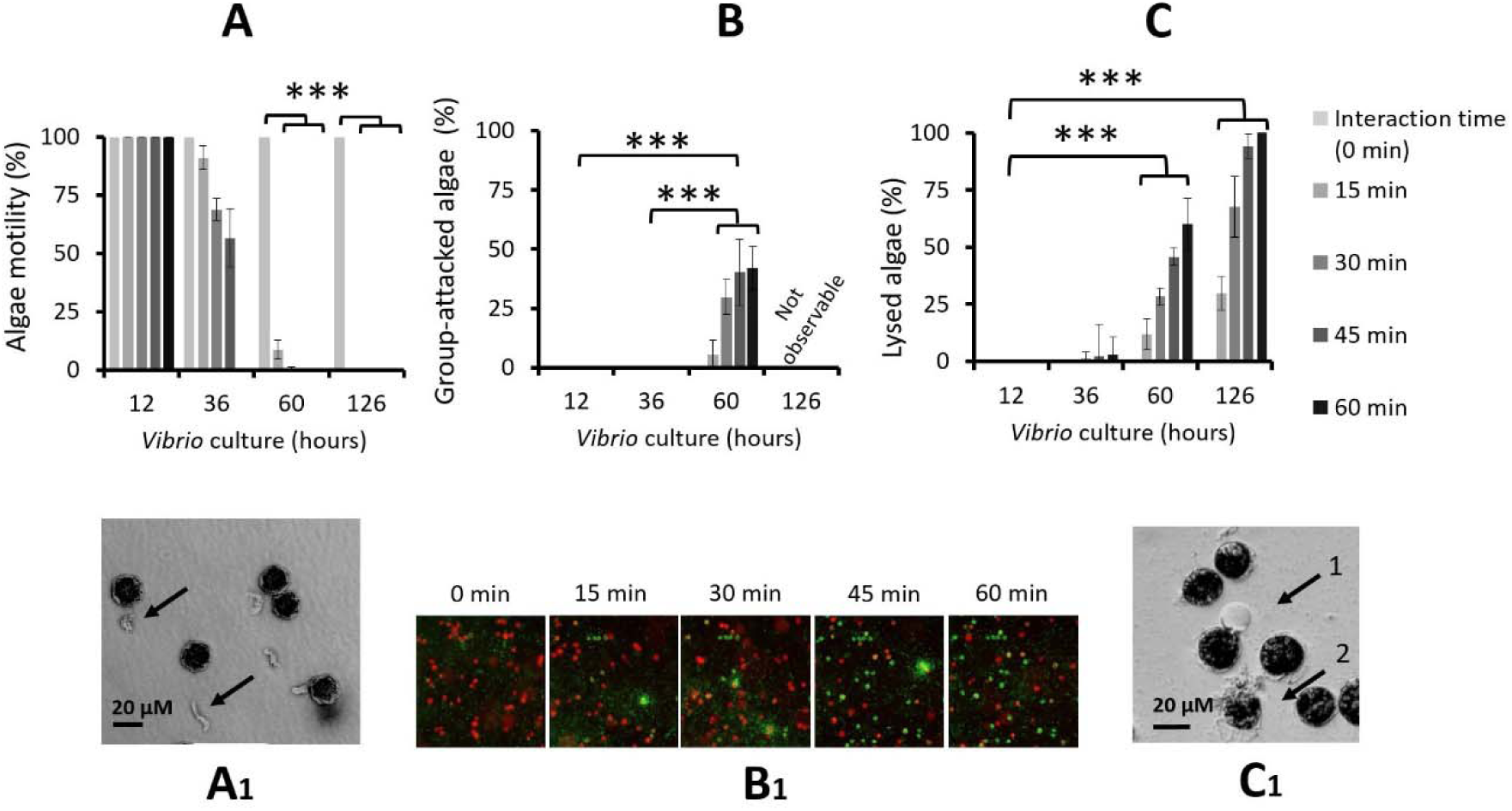
Role of *V. atlanticus* LGP32 starvation in the interspecific interaction process. Experiments were conducted by incubating *A. pacificum* ACT03 with *V. atlanticus* LGP32 previously grown for 12, 36, 60 and 126 h in Zobell medium. **(A)** Cumulative percentage of motile *A. pacificum* ACT03 cells. **(B)** Cumulative number of cells attacked by *V. atlanticus* LGP32 and **(C**) Cumulative cell lysis after 0, 15, 30, 45 and 60 minutes of interaction. Corresponding pictures showing **(A1)** Black arrows indicate unhooked and degraded flagellum from *A. pacificum* ACT03 flagellum, **(B1)** Chronological sequence of five snapshots showing *V. atlanticus* LGP32-GFP cells (60-hour culture) and *A. pacificum* ACT03 cells, during the first hour of their interaction. *V. atlanticus* LGP32 (small green cells), living *A. pacificum* ACT03 (large red cells) and dead *A. pacificum* ACT03 (large green cell). **(C1)** Black arrow 1 indicates vesicle formation on *A. pacificum* ACT03 cell and black arrow 2 indicates lysed *A. pacificum* ACT03 cell. All percentages were determined based on a minimum of 2,000 cells of *A. pacificum* ACT03. All experiments were done in triplicate. Asterisks indicate significant differences in a multiple comparison test (One-way ANOVA with post hoc Tukey test), ***P ≤ 0.001

We next tested whether this lytic effect was linked to Vibrio culture supernatant and mediated by thermostable molecule(s) secreted by Vibrio. The culture supernatant of starved culture of *V. atlanticus* LGP32 (36 h) filtered through a 10 kDa membrane and then incubated at 100°C for 30 min still possessed its lytic properties, indicating that the algicidal compounds produced by *V. atlanticus* LGP32 are small thermostable molecules unlikely to be lytic enzymes, or lysins able to digest the algae cell. However, these experimental observations clearly show the key role of nutrient limitation in triggering the attack behaviour and the secretion of lytic compounds of *V. atlanticus* LGP32.

### Attack occurs on *A. pacificum* ACT03 in exponential phase of growth

Here, we wondered whether the live/dead status of algae is important for *V. atlanticus* LGP32-mediated attacks targeting. To this end, *A. pacificum* ACT03 in exponential growth phase was first exposed for 30 minutes to the supernatant of a 126-hour culture of *V. atlanticus* LGP32, which induced lysis of 70% of the *A. pacificum* ACT03 cells (Figures 3C and 3C1 (arrow 2) and Video 4). Next, cells of *V. atlanticus* LGP32 from a 60-hour culture, capable of attacking *A. pacificum* ACT03 cells (Fig. 3B), were added. For 1 hour of exposure, no attack was observed on the previously lysed algae. This result is similar to what is observed on the Video 1 with flash attacks only on immobilized, but not degraded *A. pacificum* ACT03 cells (red), and not on lysed cells (green). In addition, no attacks occurred on cells from an old *A. pacificum* ACT03 culture (1-month culture). Together, with the very short duration of attacks (Video 1), these results indicate that *V. atlanticus* LGP32 attacked exponentially growing cells of *A. pacificum* ACT03, but not decomposing algae, suggesting that this behaviour is not just an opportunistic response of heterotrophic bacteria to organic substrates.

### Attack is independent of quorum sensing

Considering the simultaneous action of *V. atlanticus* LGP32 in attacks on *A. pacificum* ACT03, we tested whether the attack process depended on the key physiological mechanism that regulates many functions in marine microbial cells, quorum sensing (QS) (Lami, 2019; Papenfort and Bassler, 2016), a type of cell-cell communication. Although QS is a cell-density-dependent mechanism, our results showed no attack from a 12 h culture of *V. atlanticus* LGP32 up to a concentration of 4 x 10^6^ Vibrio mL^-1^ (Fig 4A). Attacks were only observed with *V. atlanticus* LGP32 from a 60 h culture at low concentration of 5 x 10^3^ Vibrio mL^-1^ to the highest concentration tested of 5 x 10^5^ Vibrio mL^-1^ (Fig 4A), consistent with the hypothesis that the attacks were independent of Vibrio density.

**Figure 4.**
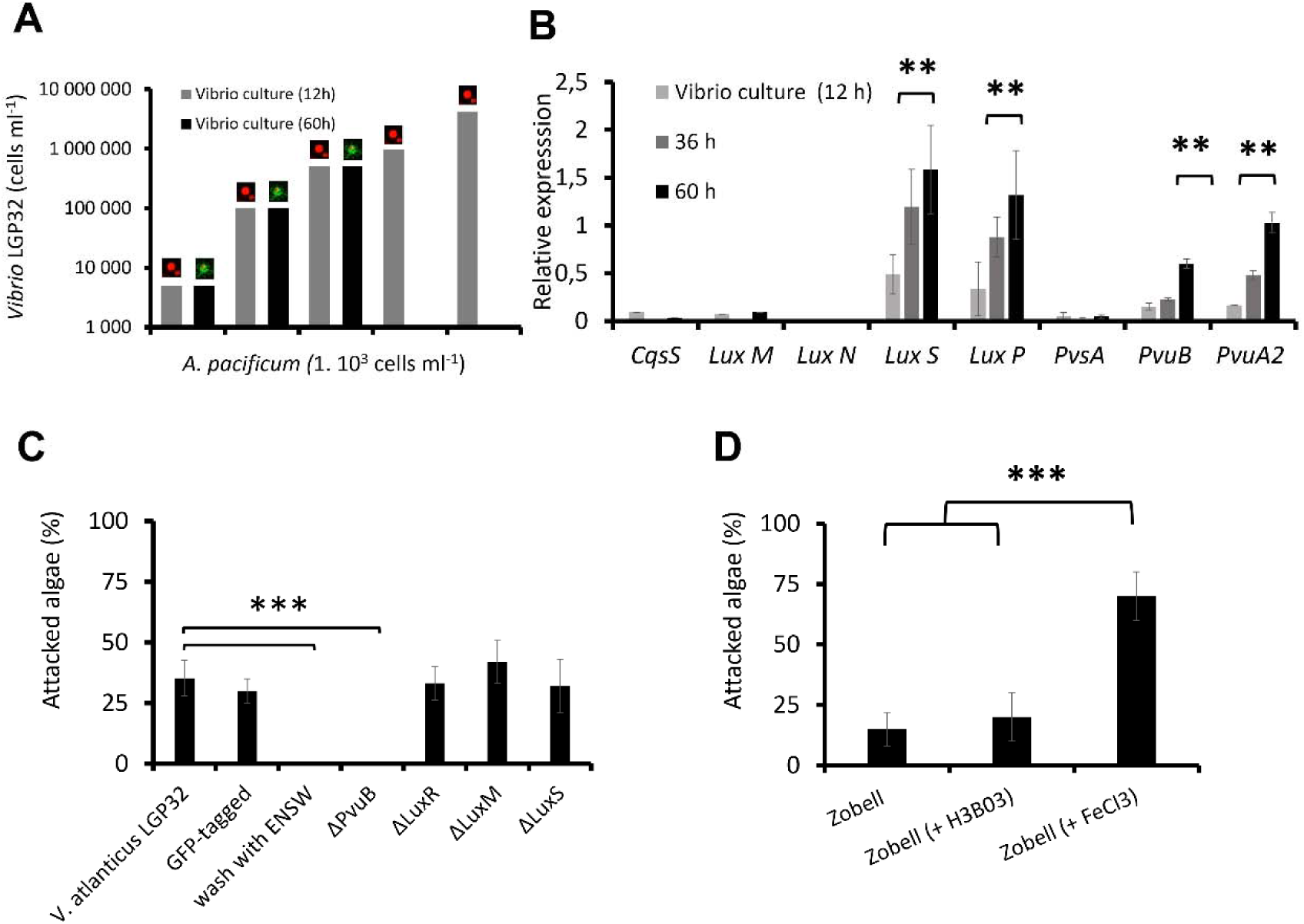
Role of quorum sensing and the vibrioferrin iron uptake pathway in the interaction process. **(A)** Effect of *V. atlanticus* LGP32 cell density on the attack process. *A. pacificum* ACT03 cells (1 x 10^3^ cells mL^-1^) were incubated with *V. atlanticus* LGP32 grown for 60 hours in Zobell medium at concentrations ranging from 5.10^3^ to 5 x 10^5^ cells mL^−1^ (black bars). For comparison, *A. pacificum* ACT03 incubated with *V. atlanticus* LGP32 grown for 12 hours in Zobell medium at concentrations ranging from 5 x 10^3^ to 4 x 10^6^ cells mL−1 (grey bars). The image on the bars indicates either unaffected algae (live red algae) or algae attacked by Vibrio (algae covered with green vibrio cell) during the interaction **(B)** CqsS, luxM, luxN, luxS, and luxP quorum sensing and PvsA, PvuB and PvuA2 vibrioferrin pathway genes expression in *V. atlanticus* LGP32 grown for 12, 36 and 60 h in Zobell medium. **(C)** Effect of *V. atlanticus* LGP32 mutants on the attack process. Experiments were conducted by incubating *A. pacificum* ACT03 with *V. atlanticus* LGP32, *V. atlanticus* LGP32 tagged with GFP, *V. atlanticus* LGP32 washed with ENSW or *V. atlanticus* LGP32 mutant ΔPvuB, ΔluxM, ΔluxR and ΔluxS previously grown 60 h in Zobell media (control), The percentage of *A. pacificum* ACT03 attacked was determined during the first 30 min of exposure. **(D)** Effect of *V. atlanticus* LGP32 culture media composition on the attack process. Experiments were conducted by incubating *A. pacificum* ACT03 with *V. atlanticus* LGP32 grown 60 h in Zobell media supplemented with boron (H_3_BO_3_) or FeCl_3._ The results were compared with an exposure to *V. atlanticus* LGP32 grown 60 h in Zobell media. All percentages were determined based on a minimum of 2,000 cells of *A. pacificum* ACT03. All experiments were done in triplicate. Asterisks indicate significant differences in a multiple comparison test (One-way ANOVA with post hoc Tukey test), **P ≤ 0.01, ***P ≤ 0.001.

The analysis of the expression of genes involved in the known QS pathways in *Vibrio* cell (Fig. S3A), highlighted that only the AI-2 pathway was induced during nutrient stress of *V. atlanticus* LGP32, because only the expression of the AI synthase (*LuxS*) and its receptor (*LuxP*) increased significantly (Fig. 4B; ANOVA p <0.05). This was confirmed by a QS bioluminescence assay, which showed AI-2 molecules (unquantified) in the Zobell culture supernatant of *V. atlanticus* LGP32 after 60 h of culture but not after 12 h of culture and not in the ENSW supernatant of *V. atlanticus* LGP32 after 12 or 60 h of culture. UHPLC-HRMS/MS provided no evidence of detectable HAI-1 and CAI-1 in any experiments.

Targeted mutagenesis of key genes involved in two of the three known QS pathways in vibrios (Fig. S3), ΔluxM (HAI-1 production), ΔluxS (AI-2 production), and ΔluxR (main high-density QS regulator), did not result in any changes in the attack behavior of *V. atlanticus* LGP32 (Fig. 4C). Combined with the absence of overexpression of the CqsS gene (inducible by CAI-1) involved in the last known QS pathway in Vibrio (Fig. S3), these results indicated that the attack by *V. atlanticus* LGP32 is most likely unrelated to QS.

### Attack related to the availability of iron

The comparative analysis of the proteome of *V. atlanticus* LGP32 incubated 60 h in artificial seawater (ENSW) versus *V. atlanticus* LGP32 grown 12 h in Zobell nutrient-rich medium revealed 10 proteins that were differentially abundant under these two contrasting conditions (Fig. S2). The two most down-regulated proteins correspond to be ß-ketoacyl-(acyl-carrier-protein) synthase II (-22-fold in ENSW compared to Zobell), a key regulator of bacterial fatty acid synthesis, and the dihydroorotase (-6.6-fold in ENSW compared to Zobell), an enzyme essential for pyrimidine biosynthesis and thus bacterial proliferation and growth. The low expression of these proteins in ENSW is consistent with *V. atlanticus* LGP32 nutritional starvation. The most up-regulated protein in starved *V. atlanticus* LGP32, with an increase of more than 6-fold, was glucosamine-6-phosphate deaminase, an enzyme involved in bacterial energy metabolism probably necessary for its survival. Among the other up-regulated proteins, one was an iron siderophore-binding protein (Spot 4413, Fig. S2A) corresponding to the vibrioferrin outer membrane receptor PvuB, whose gene is part of the *pvu* operons involved in iron transport (Fig. S3B). Interestingly, the corresponding gene *pvuB* as well as the vibrioferrin membrane receptor gene *pvuA2* (Fig. S3B) were both significantly induced in Vibrio under nutrient stress (Fig. 4B; ANOVA p <0.01) but not the one involved in the vibrioferrin biosynthesis, *pvsA* (Fig. 4B). Remarkably, of the 10 proteins identified by proteomic analysis and eliminated by mutation, only elimination of PvuB prevented *V. atlanticus* from attacking *A. pacificum* ACT03 (Fig. 4C; ANOVA p <0.001). In the absence of the *pvuB* gene, *V. atlanticus* LGP32 was unable to simultaneously attack *A. pacificum* ACT03. In addition, *V. atlanticus* LGP32 cells that had been washed with ENSW to remove their culture supernatant metabolites also failed to attack *A. pacificum* ACT03 (Fig. 4C; ANOVA p <0.001), which is congruent with the hypothesis that attacks depend on the *V. atlanticus* LGP32 vibrioferrin transport system. Finally, attacks increased significantly when FeCl_3_ was added to the Vibrio culture medium (Fig. 4D) but not with boron known to be capable of being transported by vibrioferrin (see Fig. 4D). Taken together, those results are consistent with the hypothesis that attacks are regulated by iron.

### Attack is a *Vibrio spp.* behaviour specific to *Alexandrium spp*

To evaluate the dinoflagellates specificity of the attack behaviour, a selection of *Vibrio spp.* was co-cultured with a selection of dinoflagellate strains commonly found in the Mediterranean Sea. The results showed that, among the *Vibrio spp*. tested (pathogenic or not) all, when under nutrient stress, were able to secrete algicidal compounds, immobilize, attack and lyse *A. pacificum* ACT03 cells (Table 1) and no link to their pathogenicity for fish of invertebrates was observed (Table 1). Among the sixteen dinoflagellates species tested, only *Alexandrium spp*. (non-toxic and paralytic shellfish toxin (PST) producers) and *Gymnodinium catenatum* (PST producer) were immobilized, attacked and lysed by *V. atlanticus* LGP32 (Table 2), but no link to PSTs was revealed (Table 2).

## DISCUSSION

Predation is a widespread mode of interaction for survival in the natural world (Finke and Denno, 2004; Sinclair et al., 2003). Predatory bacteria are found in a wide variety of environments and are commonly described as feeding on other bacteria, although some cases of predation on microbial eukaryotes have also been hypothesized (Johnke et al., 2014; Perez et al., 2016). Conceiving predators as free-living organisms that kill other organisms and feed on them, this study suggests that Vibrio engage in a novel form of predation in which they kill and feed on algae.

In fact, the strategy developed by Vibrio to kill algae may be reminiscent of strategies previously described in the prokaryotes (Johnke et al., 2014). As shown in Video 1, the interaction between *V. atlanticus* LGP32 and *A. pacificum* ACT03 proceeds in three stages (Fig. 5). The first stage, the ‘**immobilization stage**’, recalls the strategy used by *Streptomyces* to immobilize its prey (Kumbhar et al., 2014) based on the secretion of algicidal metabolites that disrupt the flagella. The second stage, the ‘attack stage’ corresponding to the physical contact between Vibrios and Alexandrium, is similar to the strategy used by *Myxococcus xanthus* and Lysobacter. These bacteria may require close proximity to their prey to cause lysis and utilize their biomass, although some can also kill prey at a distance through diffusible secretions ( Martin, 2002; Genovesi et al., 2013; Perez et al., 2016; Zhang et al., 2020). *V. atlanticus* LGP32 also surrounds *A. pacificum* ACT03 cells at high density for a very short time, but does not invade the algal cell. Visually, this phenomenon resembles bacteria clustering around lysed ciliate cells (Blackburn et al., 1998). The third stage, the ‘**killing stage**’, is similar to that of epibiotic bacterial predators, which induce the lysis of bacteria from the outside (Rashidan and Bird, 2001). Overall, these observations suggest that *V. atlanticus* LGP32 can exhibit a predatory behavior.

**Figure 5.**
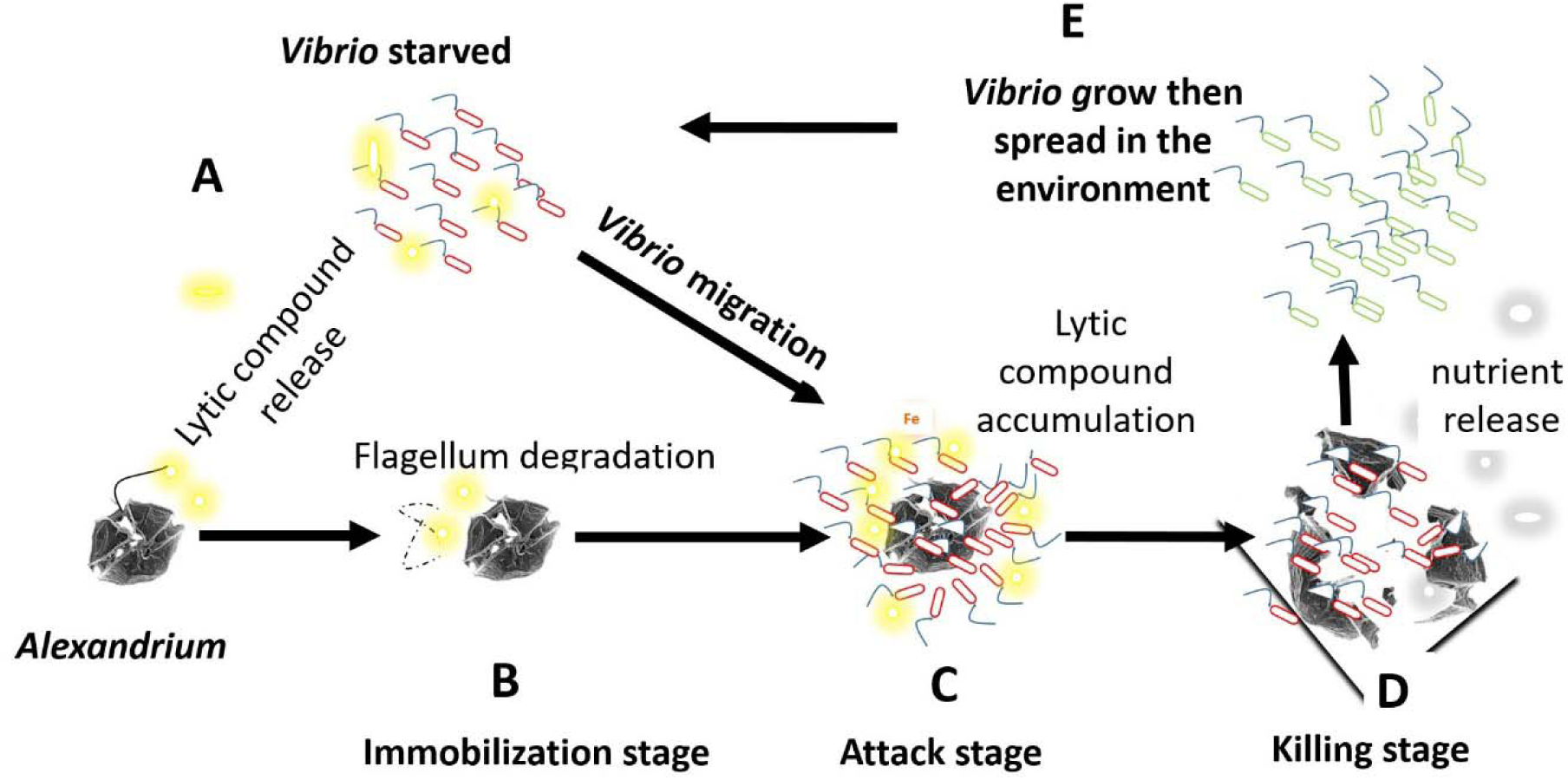
Schematic representation of a putative strategy developed by *Vibrio spp*. to feed on Alexandrium *spp*. and *G. catenatum* in the environment. **(A)** Vibrio in the environment when subjected to starvation secrete non-protein lytic compounds. **(B)** Some of these lytic compounds degrade the flagella, immobilizing the alga (immobilization stage). **(C)** Then Vibrio swims and clusters around its prey (attack stage). **(D)** Lytic compounds released by Vibrio where able to concentrate around the algal cells, thereby lysing the algae (killing stage). **(E)** Feeding on the released nutrients, Vibrio multiply and then spread in the environment. Yellow clouds: Lytic compound release by Vibrio, Grey clouds: Algal nutrients released upon lysis.

The attack behaviour of *V. atlanticus* LGP32 was linked to iron absorption, as mutants with impaired iron absorption completely lost its ability to attack the algae. Iron is an essential element for growth in most organisms, including phytoplankton (Martin and Fitzwater, 1988) and bacteria (Neilands, 1981), and its concentration in seawater is known to be very low with measurements in the open ocean surface at 0.2 nM and in deep waters at 0.6 nM (Millero, 1998).

Moreover, its low solubility in seawater limits its availability (Bruland et al., 1994; Wu and Luther, 1994). To acquire iron, bacteria have developed systems based on the secretion (and subsequent uptake) of iron-chelating siderophores to obtain this element from the environment (Amin et al., 2009a). Therefore, many *Vibrio spp*. produce a siderophore known as vibrioferrin, which is synthesized and secreted by proteins encoded by the *pvsABCDE* gene cluster (Fig. S3). Given that boron is known for its role in regulating a global bacterial cellular response to phytoplankton and to bind to vibrioferrin (Romano et al., 2013; Weerasinghe et al., 2013), we tested its potential involvement in simultaneous Vibrio attacks. Compared to the Zobell control, no effect on the number of attacks was observed. For iron-vibrioferrin uptake, *Vibrio parahaemolyticus* uses a membrane siderophore receptor, called PvuA, which is coupled to an inner membrane ATP-binding cassette (ABC). This ABC transporter system, which is comprised of proteins encoded by the pvuABCDE gene cluster (Fig. S3), is required for the transport of the siderophore across the inner membrane (Tanabe et al., 2003). Siderophores are not only iron carriers but also important regulators of virulence (Miethke and Marahiel, 2007) and mediators of bacterial interaction with phytoplankton (Amin et al., 2009b; Kramer et al., 2020). We showed here a pivotal role of iron in the interaction between *V. atlanticus* LGP32 and *A. pacificum* ACT03. This mirrors the mutualistic interaction observed between *Gymnodinium catenatum* and *Marinobacter* (Amin et al., 2009b). In fact, in natural settings, the co-occurrence of *Marinobacter* and *G. catenatum* is suggested to depend on a mutually beneficial utilization of iron and carbon resources (Bolch et al., 2011). As in the present study, iron seems to play a key role in the interaction. Indeed, the labile iron released through the photolysis of ferric chelates with vibrioferrin providing a crucial iron source for phytoplankton, which need substantial amounts of iron to support carbon fixation through photosynthesis (Amin et al., 2009a; Yang et al., 2021). This fixed carbon, in turn, sustains the growth of both the phytoplankton and their associated bacterial counterparts (Amin et al., 2009b; Kramer et al., 2020). Interestingly, if a general nutrient deficiency causes attacks, iron supplementation increases this number of attacks (Figure 4D), suggesting the importance of iron absorption in the attack behavior. Future studies should determine whether a nutrient deficiency increases the iron absorption capacity of Vibrio bacteria and whether this could play a major role in the attack mechanism.

This study showed that quorum sensing is not involved in microbial attacks. Thus, *V. atlanticus* LGP32 mutants lacking the genes involved in known quorum sensing pathways exhibit the same phenotype as wild-type *V. atlanticus* LGP32, and Vibrio density does not induce attacks. However, V. atlanticus LGP32 produces AI-2 during attacks. Quorum sensing and iron acquisition are sometimes interconnected in Vibrio (McRose et al., 2018). For example, in *Vibrio vulnificus*, the production of vulnibactin (a siderophore) is known to be controlled by AI-2 (Kim and Shin, 2011). Similarly, AI-2 could be involved in the production of vibrioferrin in *V. atlanticus* LGP32.

In the natural environment, associations between bacteria and algae have already been observed (Lopez-Joven et al., 2018; Miller et al., 2005; Rosales et al., 2022; Xu et al., 2022). We have shown here that algal attacks by Vibrio can be carried out *in vitro*. Environmental data collected in the Thau lagoon showed a correlation between the presence of Alexandrium and that of Vibrio, suggesting that such interactions could also occur in the marine environment. If that were the case, this behaviour would provide an important ecological advantage to Vibrio to obtain nutrients in the environment, where *Alexandrium spp*. and *Gymnodinium catenatum* form blooms.

With more than 30 species distributed all over the world (Anderson et al., 2012; Hallegraeff et al., 2021), *Alexandrium spp.* and *Gymnodinium spp*. considered as invasive species by the Delivering Alien Invasive Species Inventories for Europe (http://www.europe-aliens.org)), could play an unexpected and important role in maintaining, structuring and regulating Vibrio populations in the ecosystem. In turn, Vibrio could contribute to the regulation and control of their blooms.

To conclude, this study reveals the capacity of some *Vibrio spp*. to act as facultative predator-like bacteria that hunt specific algae. In the current context of climate change, which is favourable to their development, monitoring the invasive algae *Alexandrium spp*. and *Gymnodinium catenatum* should be considered not only for their potent harmful effect on humans and animals, but also because they may represent a potential source of nutrients for the expansion of Vibrio, particularly pathogenic species (Lemire et al., 2015).

## DATA AVAILABILITY

All data on environmental study are included in the article and/or supporting information files. All other study data are available on request from the corresponding author.

## ACKNOWLEDGMENT

The authors thank F. Le Roux and Y. Labreuche for their gift of *Vibrio* mutants; E. Garcés, S. Turki, H. Frehi, A. Bouquet and W. Medhioub for their contribution in obtaining Mediterranean phytoplankton strains; K. Escoubeyrou and D. Stien for advice on HRMS/MS data; V. Diakou-Verdin and E. Jublanc from the Montpellier RIO Imaging Platform for access to microscope facilities (www.mri.cnrs.fr); P. Clair from the Montpellier qPHD platform and the Technoviv platform of the UPVD for access to PCR facilities; Luc Markiw, N. Brunet, A. Lang, A. Payelleville, T. Milhau and M. Leroy for their technical assistance; The IFREMER LER-LR laboratory for access to the Thau Lagoon; the BIO2MAR platform at the Observatoire Océanologique de Banyuls for providing access to UHPLC-HRMS/MS facilities.

## FUNDING

European VIVALDI grant 678589, French EC2CO ROSEOCOM grant and IFREMER ALGOVIR grant.

## FIGURES, TABLES and VIDEOS

**Figure S1.**
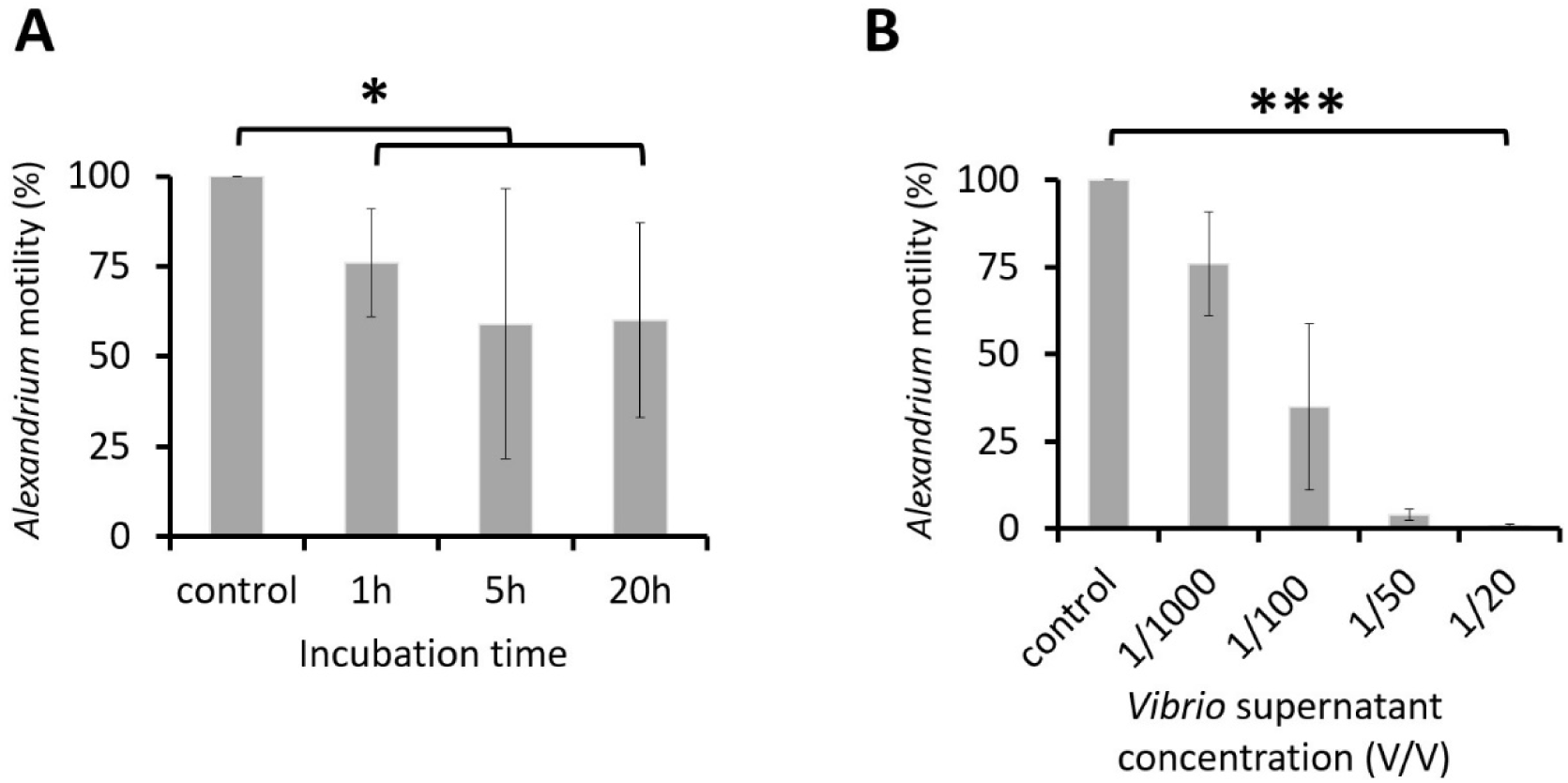
Time and dose-dependent effects of the *V. atlanticus* LGP32 culture supernatant on *A. pacificum* ACT03 motility. **(A)** A time dependence experiment was conducted by incubating *A. pacificum* ACT03 for 1, 5 or 20 h with 1/1000 v/v (1 µL/mL) of culture supernatant from *V. atlanticus* LGP32 previously grown for 60 h in Zobell culture media. **(B)** A dose-dependence experiment was conducted by incubating *A. pacificum* ACT03 for 1 h with 1/1000 to 1/20 v/v (1-50 µL/mL) of culture supernatant from *V. atlanticus* previously grown for 60 h in Zobell media. The percentage of motile *A. pacificum* ACT03 was determined after 1 hour of exposure. All percentages were determined based on a minimum of 2,000 cells of *A. pacificum* ACT03. Error bars represent the standard deviation of the mean of three independent experiments. Asterisks indicate significant differences in a multiple comparison test (One-way ANOVA with post hoc Tukey test), *P ≤ 0.05, ***P ≤ 0.001.

**Figure S2.**
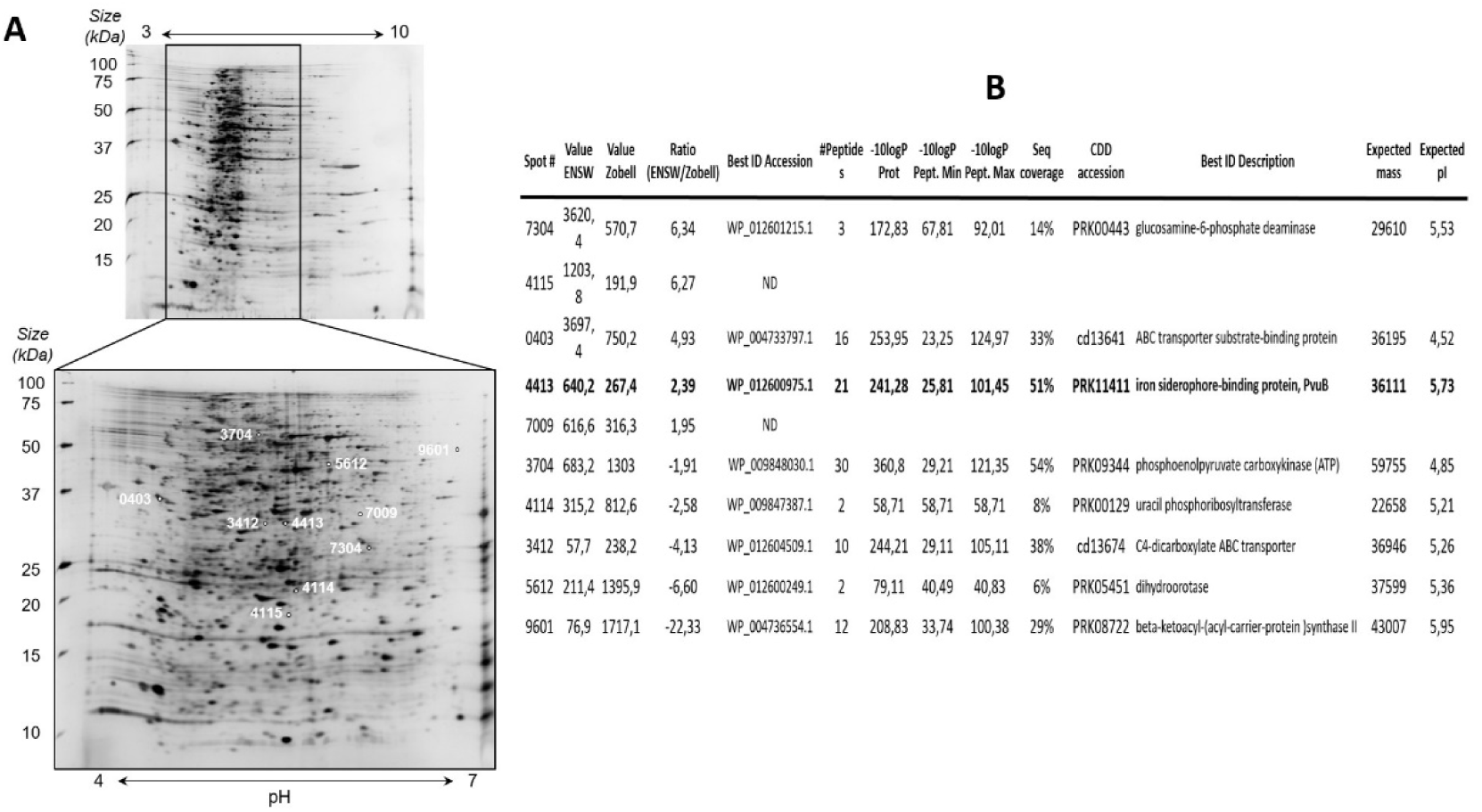
*V. atlanticus* LGP32 proteome analysis following nutrient stress. **(A)** Example of 2D gel, the numbers in white on the gel 4-7 correspond to the number and position of the protein spots analyzed. **(B)** Proteins identified by LC-MS/MS as differentially represented in the 2D gel comparative approach following nutrient stress. ND: Not determined; ENSW (artificial seawater).

**Figure S3.**
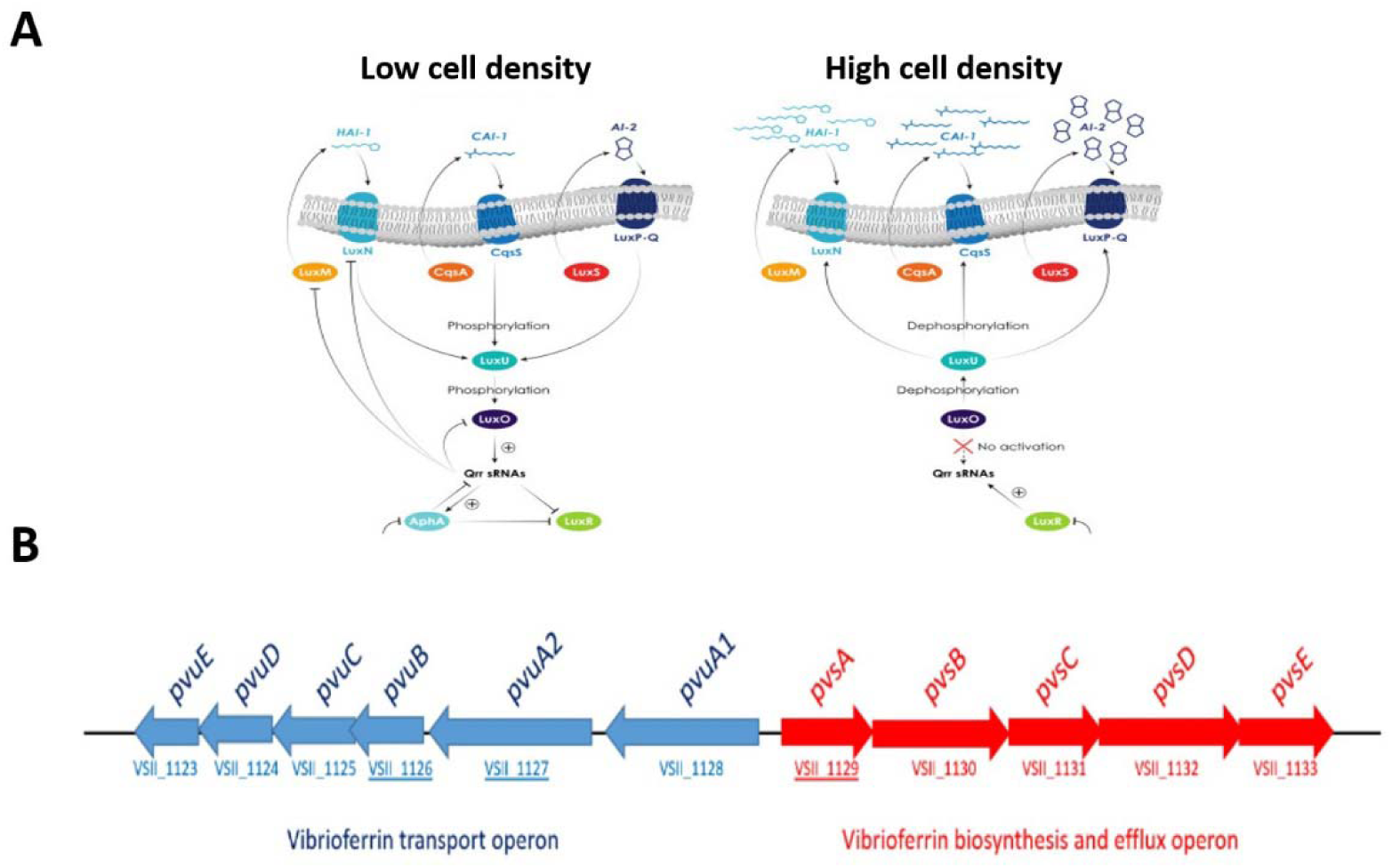
Quorum sensing and the vibrioferrin iron uptake pathway in Vibrio. **(A)** Putative quorum sensing (QS) pathways at low and high cell density in Vibrio according to Lami et al.(Lami, 2019). **(B)** Genetic organization of the vibrioferrin utilization gene cluster on *V. atlanticus* LGP32 chromosome 2. The Pvu and Pvs operons are involved in the secretion and the transport of ferric vibrioferrin and biosynthesis of vibrioferrin, respectively. Arrows indicate the transcriptional directions of the genes. VSII1126, VSII1137 and VSII1129 corresponding to PvuB, PvuA2 and PvsA genes respectively.

## VIDEOS

**Video 1. Dynamics of *V. atlanticus* LGP32*-Alexandrium pacificum* ACT03 interaction.** GFP-tagged *V. atlanticus* (small green cells); living *A. pacificum* (large red cells); lysed *A. pacificum* (large green cells) filmed under an epifluorescence microscope.

**Video 2. Second-by-second timing of *V. atlanticus* LGP32 attacking *Alexandrium pacificum* ACT03.** GFP-tagged *V. atlanticus* (small green cells); *A. pacificum* living cell (large red cells) filmed under an epifluorescence microscope.

**Video 3. Degradation and disruption of *Alexandrium pacificum* ACT03 flagella.** Effect of Vibrio supernatant on the first stage of the interaction filmed under a confocal microscope.

**Video 4. Attacks of *V. atlanticus* LGP32 on target *Alexandrium pacificum* ACT03.** This video, recorded under a confocal microscope, shows Vibrios simultaneously attacking a first immobilized Alexandrium cell, then moving on to attack a second cell without ever targeting the other cells present, suggesting active communication between the Vibrio bacteria. *V. atlanticus* LGP32 (small cells); *A. pacificum* ACT03 (large cells).

**Video 5. Vesicle formation and bursting of an *Alexandrium pacificum* ACT03.** Direct effect of Vibrio supernatant on *Alexandrium* after 126 h of culture filmed under a confocal microscope.

## Notes

### Competing Interest Statement

The authors have declared no competing interest.

### Summary of Updates

This revised version includes updates to the abstract, main text, figures and tables.

## REFERENCES

Abadie, E., Amzil, Z., Belin, C., Comps, M.A., Elzière-Papayanni, P., Lassus, P., le Bec, C., Baut, M.L., Nezan, E., Poggi, R., 1999. Contamination de l’étang de Thau par Alexandrium tamarense (No. Rapport 884). Ifremer.

Abadie, E., Kaci, L., Berteaux, T., Hess, P., Sechet, V., Masseret, E., Rolland, J.L., Laabir, M., 2015. Effect of Nitrate, Ammonium and Urea on Growth and Pinnatoxin G Production of Vulcanodinium rugosum. Mar. Drugs 13, 5642–5656. 10.3390/md13095642

Amin, S.A., Green, D.H., Hart, M.C., Kuepper, F.C., Sunda, W.G., Carrano, C.J., 2009a. Photolysis of iron-siderophore chelates promotes bacterial-algal mutualism. Proc. Natl. Acad. Sci. U. S. A. 106, 17071–17076. 10.1073/pnas.0905512106

Amin, S.A., Green, D.H., Kuepper, F.C., Carrano, C.J., 2009b. Vibrioferrin, an Unusual Marine Siderophore: Iron Binding, Photochemistry, and Biological Implications. Inorg. Chem. 48, 11451–11458. 10.1021/ic9016883

Anderson, D.M., Alpermann, T.J., Cembella, A.D., Collos, Y., Masseret, E., Montresor, M., 2012. The globally distributed genus Alexandrium: Multifaceted roles in marine ecosystems and impacts on human health. Harmful Algae 14, 10–35. 10.1016/j.hal.2011.10.012

Baker-Austin, C., Trinanes, J., Gonzalez-Escalona, N., Martinez-Urtaza, J., 2017. Non-Cholera Vibrios: The Microbial Barometer of Climate Change. Trends Microbiol. 25, 76–84. 10.1016/j.tim.2016.09.008

Ben-Gharbia, H., Yahia, O.K.-D., Amzil, Z., Chomerat, N., Abadie, E., Masseret, E., Sibat, M., Triki, H.Z., Nouri, H., Laabir, M., 2016. Toxicity and Growth Assessments of Three Thermophilic Benthic Dinoflagellates (Ostreopsis cf. ovata, Prorocentrum lima and Coolia monotis) Developing in the Southern Mediterranean Basin. Toxins 8, 297. 10.3390/toxins8100297

Blackburn, N., Fenchel, T., Mitchell, J., 1998. Microscale nutrient patches in planktonic habitats shown by chemotactic bacteria. Science 282, 2254–2256. 10.1126/science.282.5397.2254

Bolch, C.J.S., Subramanian, T.A., Green, D.H., 2011. THE TOXIC DINOFLAGELLATE GYMNODINIUM CATENATUM (DINOPHYCEAE) REQUIRES MARINE BACTERIA FOR GROWTH^1^. J. Phycol. 47, 1009–1022. 10.1111/j.1529-8817.2011.01043.x

Bruland, K., Orians, K., Cowen, J., 1994. Reactive Trace-Metals in the Stratified Central North Pacific. Geochim. Cosmochim. Acta 58, 3171–3182. 10.1016/0016-7037(94)90044-2

Brumfield, K.D., Usmani, M., Chen, K.M., Gangwar, M., Jutla, A.S., Huq, A., Colwell, R.R., 2021. Environmental parameters associated with incidence and transmission of pathogenic Vibrio spp. Environ. Microbiol. 23, 7314–7340. 10.1111/1462-2920.15716

Burkholder, J.M., Shumway, S.E., Glibert, P.M., 2018. Food Web and Ecosystem Impacts of Harmful Algae, in: Harmful Algal Blooms. John Wiley & Sons, Ltd, pp. 243–336. 10.1002/9781118994672.ch7

Burnham, K.P., Anderson, D.R. (Eds.), 2002. Summary, in: Model Selection and Multimodel Inference: A Practical Information-Theoretic Approach. Springer, New York, NY, pp. 437–454. 10.1007/978-0-387-22456-5_8

Chai, Z.Y., Wang, H., Deng, Y., Hu, Z., Tang, Y.Z., 2020. Harmful algal blooms significantly reduce the resource use efficiency in a coastal plankton community. Sci. Total Environ. 704, 135381. 10.1016/j.scitotenv.2019.135381

Coyne, K.J., Wang, Y., Johnson, G., 2022. Algicidal Bacteria: A Review of Current Knowledge and Applications to Control Harmful Algal Blooms. Front. Microbiol. 13, 871177. 10.3389/fmicb.2022.871177

Drebes Dörr, N.C., Blokesch, M., 2020. Interbacterial competition and anti-predatory behaviour of environmental Vibrio cholerae strains. Environ. Microbiol. 22, 4485–4504. 10.1111/1462-2920.15224

Espinoza-Vergara, G., Hoque, M.M., McDougald, D., Noorian, P., 2020. The Impact of Protozoan Predation on the Pathogenicity of Vibrio cholerae. Front. Microbiol. 11, 17. 10.3389/fmicb.2020.00017

Fertouna-Bellakhal, M., Dhib, A., Fathalli, A., Bellakhal, M., Chomérat, N., Masseret, E., Laabir, M., Turki, S., Aleya, L., 2015. Alexandrium pacificum Litaker sp. nov (Group IV): Resting cyst distribution and toxin profile of vegetative cells in Bizerte Lagoon (Tunisia, Southern Mediterranean Sea). Harmful Algae 48, 69–82. 10.1016/j.hal.2015.07.007

Finke, D.L., Denno, R.F., 2004. Predator diversity dampens trophic cascades. Nature 429, 407–410. 10.1038/nature02554

Gay, M., Renault, T., Pons, A.M., Le Roux, F., 2004. Two Vibrio splendidus related strains collaborate to kill Crassostrea gigas: taxonomy and host alterations. Dis. Aquat. Organ. 62, 65–74. 10.3354/dao062065

Genovesi, B., Mouillot, D., Laugier, T., Fiandrino, A., Laabir, M., Vaquer, A., Grzebyk, D., 2013. Influences of sedimentation and hydrodynamics on the spatial distribution of Alexandrium catenella/tamarense resting cysts in a shellfish farming lagoon impacted by toxic blooms. Harmful Algae 25, 15–25. 10.1016/j.hal.2013.02.002

Genovesi, B., Shin-Grzebyk, M.-S., Grzebyk, D., Laabir, M., Gagnaire, P.-A., Vaquer, A., Pastoureaud, A., Lasserre, B., Collos, Y., Berrebi, P., Masseret, E., 2011. Assessment of cryptic species diversity within blooms and cyst bank of the Alexandrium tamarense complex (Dinophyceae) in a Mediterranean lagoon facilitated by semi-multiplex PCR. J. Plankton Res. 33, 405–414. 10.1093/plankt/fbq127

Hadjadji, I., Laabir, M., Frihi, H., Collos, Y., Shao, Z.J., Berrebi, P., Abadie, E., Amzil, Z., Chomérat, N., Rolland, J.L., Rieuvilleneuve, F., Masseret, E., 2020. Unsuspected intraspecific variability in the toxin production, growth and morphology of the dinoflagellate Alexandrium pacificum R.W. Litaker (Group IV) blooming in a South Western Mediterranean marine ecosystem, Annaba Bay (Algeria). Toxicon 180, 79–88. 10.1016/j.toxicon.2020.04.005

Hallegraeff, G.M., Anderson, D.M., Belin, C., Bottein, M.-Y.D., Bresnan, E., Chinain, M., Enevoldsen, H., Iwataki, M., Karlson, B., McKenzie, C.H., Sunesen, I., Pitcher, G.C., Provoost, P., Richardson, A., Schweibold, L., Tester, P.A., Trainer, V.L., Yñiguez, A.T., Zingone, A., 2021. Perceived global increase in algal blooms is attributable to intensified monitoring and emerging bloom impacts. Commun. Earth Environ. 2, 117. 10.1038/s43247-021-00178-8

Harrison, P., Waters, R., Taylor, F., 1980. A Broad-Spectrum Artificial Seawater Medium for Coastal and Open Ocean Phytoplankton. J. Phycol. 16, 28–35. 10.1111/j.1529-8817.1980.tb00724.x

Johnke, J., Cohen, Y., de Leeuw, M., Kushmaro, A., Jurkevitch, E., Chatzinotas, A., 2014. Multiple micro-predators controlling bacterial communities in the environment. Curr. Opin. Biotechnol. 27, 185–190. 10.1016/j.copbio.2014.02.003

Johnson, C.N., 2013. Fitness Factors in Vibrios: a Mini-review. Microb. Ecol. 65, 826–851. 10.1007/s00248-012-0168-x

Kim, C.M., Shin, S.H., 2011. Modulation of Iron-Uptake Systems by a Mutation of luxS Encoding an Autoinducer-2 Synthase in Vibrio vulnificus. Biol. Pharm. Bull. 34, 632–637. 10.1248/bpb.34.632

Kitatsukamoto, K., Oyaizu, H., Nanba, K., Simidu, U., 1993. Phylogenetic-Relationships of Marine-Bacteria, Mainly Members of the Family Vibrionaceae, Determined on the Basis of 16s Ribosomal-Rna Sequences. Int. J. Syst. Bacteriol. 43, 8–19. 10.1099/00207713-43-1-8

Kramer, J., Oezkaya, O., Kuemmerli, R., 2020. Bacterial siderophores in community and host interactions. Nat. Rev. Microbiol. 18, 152–163. 10.1038/s41579-019-0284-4

Kumbhar, C., Mudliar, P., Bhatia, L., Kshirsagar, A., Watve, M., 2014. Widespread predatory abilities in the genus Streptomyces. Arch. Microbiol. 196, 235–248. 10.1007/s00203-014-0961-7

Laabir, M., Jauzein, C., Genovesi, B., Masseret, E., Grzebyk, D., Cecchi, P., Vaquer, A., Perrin, Y., Collos, Y., 2011. Influence of temperature, salinity and irradiance on the growth and cell yield of the harmful red tide dinoflagellate Alexandrium catenella colonizing Mediterranean waters. J. Plankton Res. 33, 1550–1563. 10.1093/plankt/fbr050

Laanaia, N., Vaquer, A., Fiandrino, A., Genovesi, B., Pastoureaud, A., Cecchi, P., Collos, Y., 2013. Wind and temperature controls on Alexandrium blooms (2000-2007) in Thau lagoon (Western Mediterranean). Harmful Algae 28, 31–36. 10.1016/j.hal.2013.05.016

Labreuche, Y., Le Roux, F., Henry, J., Zatylny, C., Huvet, A., Lambert, C., Soudant, P., Mazel, D., Nicolas, J.-L., 2010. Vibrio aestuarianus zinc metalloprotease causes lethality in the Pacific oyster Crassostrea gigas and impairs the host cellular immune defenses. Fish Shellfish Immunol. 29, 753–758. 10.1016/j.fsi.2010.07.007

Lami, R., 2019. Quorum Sensing in Marine Biofilms and Environments, in: Quorum Sensing, Academic Press. Elsevier, pp. 55–96. 10.1016/B978-0-12-814905-8.00003-4

Le Roux, F., Binesse, J., Saulnier, D., Mazel, D., 2007. Construction of a Vibrio splendidus Mutant Lacking the Metalloprotease Gene vsm by Use of a Novel Counterselectable Suicide Vector. Appl. Environ. Microbiol. 73, 777–784. 10.1128/AEM.02147-06

Leblad, B.R., Amnhir, R., Reqia, S., Sitel, F., Daoudi, M., Marhraoui, M., Abdellah, M.K.O., Veron, B., Er-Raioui, H., Laabir, M., 2020. Seasonal variations of phytoplankton assemblages in relation to environmental factors in Mediterranean coastal waters of Morocco, a focus on HABs species. Harmful Algae 96, 101819. 10.1016/j.hal.2020.101819

Lemire, A., Goudenege, D., Versigny, T., Petton, B., Calteau, A., Labreuche, Y., Le Roux, F., 2015. Populations, not clones, are the unit of vibrio pathogenesis in naturally infected oysters. Isme J. 9, 1523–1531. 10.1038/ismej.2014.233

LeRoux, F., Wegner, K.M., Baker-Austin, C., Vezzulli, L., Osorio, C.R., Amaro, C., Ritchie, J.M., Defoirdt, T., Destoumieux-Garzon, D., Blokesch, M., Mazel, D., Jacq, A., Cava, F., Gram, L., Wendling, C.C., Strauch, E., Kirschner, A., Huehn, S., 2015. The emergence of Vibrio pathogens in Europe: ecology, evolution, and pathogenesis (Paris, 11-12th March 2015). Front. Microbiol. 6, 830. 10.3389/fmicb.2015.00830

Li, D., Zhang, H., Fu, L., An, X., Zhang, B., Li, Y., Chen, Z., Zheng, W., Yi, L., Zheng, T., 2014. A Novel Algicide: Evidence of the Effect of a Fatty Acid Compound from the Marine Bacterium, Vibrio sp BS02 on the Harmful Dinoflagellate, Alexandrium tamarense. Plos One 9, e91201. 10.1371/journal.pone.0091201

Liu, P.-C., Lee, K.-K., Yii, K.-C., Kou, G.-H., Chen, S.-N., 1996. News & Notes: Isolation of Vibrio harveyi from Diseased Kuruma Prawns Penaeus japonicus. Curr. Microbiol. 33, 129–132. 10.1007/s002849900087

Lopez-Joven, C., Rolland, J.-L., Haffner, P., Caro, A., Roques, C., Carre, C., Travers, M.-A., Abadie, E., Laabir, M., Bonnet, D., Destoumieux-Garzon, D., 2018. Oyster Farming, Temperature, and Plankton Influence the Dynamics of Pathogenic Vibrios in the Thau Lagoon. Front. Microbiol. 9, 2530. 10.3389/fmicb.2018.02530

Mandel, M.J., Stabb, E.V., Ruby, E.G., 2008. Comparative genomics-based investigation of resequencing targets in Vibrio fischeri: Focus on point miscalls and artefactual expansions. Bmc Genomics 9, 138. 10.1186/1471-2164-9-138

Marampouti, C., Buma, A.G.J., De Boer, M.K., 2021. Mediterranean alien harmful algal blooms: origins and impacts. Environ. Sci. Pollut. Res. 28, 3837–3851. 10.1007/s11356-020-10383-1

Martin, J., Fitzwater, S., 1988. Iron-Deficiency Limits Phytoplankton Growth in the Northeast Pacific Subarctic. Nature 331, 341–343. 10.1038/331341a0

Martin, M.O., 2002. Predatory prokaryotes: An emerging research opportunity. J. Mol. Microbiol. Biotechnol. 4, 467–477.

Mavian, C., Paisie, T.K., Alam, M.T., Browne, C., Beau De Rochars, V.M., Nembrini, S., Cash, M.N., Nelson, E.J., Azarian, T., Ali, A., Morris, J.G., Salemi, M., 2020. Toxigenic Vibrio cholerae evolution and establishment of reservoirs in aquatic ecosystems. Proc. Natl. Acad. Sci. 117, 7897–7904. 10.1073/pnas.1918763117

McRose, D.L., Baars, O., Seyedsayamdost, M.R., Morel, F.M.M., 2018. Quorum sensing and iron regulate a two-for-one siderophore gene cluster in Vibrio harveyi. Proc. Natl. Acad. Sci. 115, 7581–7586. 10.1073/pnas.1805791115

Miethke, M., Marahiel, M.A., 2007. Siderophore-Based Iron Acquisition and Pathogen Control. Microbiol. Mol. Biol. Rev. 71, 413–451. 10.1128/MMBR.00012-07

Miller, S.D., Haddock, S.H.D., Elvidge, C.D., Lee, T.F., 2005. Detection of a bioluminescent milky sea from space. Proc. Natl. Acad. Sci. U. S. A. 102, 14181–14184. 10.1073/pnas.0507253102

Millero, F.J., 1998. Solubility of Fe(III) in seawater. Earth Planet. Sci. Lett. 154, 323–329. 10.1016/S0012-821X(97)00179-9

Muhling, B.A., Jacobs, J., Stock, C.A., Gaitan, C.F., Saba, V.S., 2017. Projections of the future occurrence, distribution, and seasonality of three Vibrio species in the Chesapeake Bay under a high-emission climate change scenario. Geohealth 1, 278–296. 10.1002/2017GH000089

Neilands, J., 1981. Iron-Absorption and Transport in Microorganisms. Annu. Rev. Nutr. 1, 27–46. 10.1146/annurev.nu.01.070181.000331

Oberbeckmann, S., Fuchs, B.M., Meiners, M., Wichels, A., Wiltshire, K.H., Gerdts, G., 2012. Seasonal Dynamics and Modeling of a Vibrio Community in Coastal Waters of the North Sea. Microb. Ecol. 63, 543–551. 10.1007/s00248-011-9990-9

Pal, M., Yesankar, P.J., Dwivedi, A., Qureshi, A., 2020. Biotic control of harmful algal blooms (HABs): A brief review. J. Environ. Manage. 268, 110687. 10.1016/j.jenvman.2020.110687

Papenfort, K., Bassler, B.L., 2016. Quorum sensing signal-response systems in Gram-negative bacteria. Nat. Rev. Microbiol. 14, 576–588. 10.1038/nrmicro.2016.89

Perez, J., Moraleda-Munoz, A., Javier Marcos-Torres, F., Munoz-Dorado, J., 2016. Bacterial predation: 75 years and counting! Environ. Microbiol. 18, 766–779. 10.1111/1462-2920.13171

Pfaffl, M.W., 2001. A new mathematical model for relative quantification in real-time RT-PCR. Nucleic Acids Res. 29, e45. 10.1093/nar/29.9.e45

R Core Team, 2024. R: A Language and Environment for Statistical Computing. https://www.R-project.org/

Rashidan, K.K., Bird, D.F., 2001. Role of predatory bacteria in the termination of a cyanobacterial bloom. Microb. Ecol. 41, 97–105.

Rodrigues, A.M.S., Lami, R., Escoubeyrou, K., Intertaglia, L., Mazurek, C., Doberva, M., Pérez-Ferrer, P., Stien, D., 2022. Straightforward N -Acyl Homoserine Lactone Discovery and Annotation by LC–MS/MS-based Molecular Networking. J. Proteome Res. 21, 635–642. 10.1021/acs.jproteome.1c00849

Rolland, J.-L., Pelletier, K., Masseret, E., Rieuvilleneuve, F., Savar, V., Santini, A., Amzil, Z., Laabir, M., 2012. Paralytic Toxins Accumulation and Tissue Expression of alpha-Amylase and Lipase Genes in the Pacific Oyster Crassostrea gigas Fed with the Neurotoxic Dinoflagellate Alexandrium catenella. Mar. Drugs 10, 2519–2534. 10.3390/md10112519

Romano, A., Trimble, L., Hobusch, A.R., Schroeder, K.J., Amin, S.A., Hartnett, A.D., Barker, R.A., Crumbliss, A.L., Carrano, C.J., 2013. Regulation of iron transport related genes by boron in the marine bacterium Marinobacter algicola DG893. Metallomics 5, 1025. 10.1039/c3mt00068k

Rosales, D., Ellett, A., Jacobs, J., Ozbay, G., Parveen, S., Pitula, J., 2022. Investigating the Relationship between Nitrate, Total Dissolved Nitrogen, and Phosphate with Abundance of Pathogenic Vibrios and Harmful Algal Blooms in Rehoboth Bay, Delaware. Appl. Environ. Microbiol. 88, e00356–22. 10.1128/aem.00356-22

Sinclair, A.R.E., Mduma, S., Brashares, J.S., 2003. Patterns of predation in a diverse predator-prey system. Nature 425, 288–290. 10.1038/nature01934

Su, R.Q., Yang, X.R., Zheng, T.L., Tian, Y., Jiao, N.Z., Cai, L.Z., Hong, H.S., 2007. Isolation and characterization of a marine algicidal bacterium against the toxic dinoflagellate Alexandrium tamarense. Harmful Algae 6, 799–810. 10.1016/j.hal.2007.04.004

Tanabe, T., Funahashi, T., Nakao, H., Miyoshi, S.-I., Shinoda, S., Yamamoto, S., 2003. Identification and Characterization of Genes Required for Biosynthesis and Transport of the Siderophore Vibrioferrin in Vibrio parahaemolyticus. J. Bacteriol. 185, 6938–6949. 10.1128/JB.185.23.6938-6949.2003

Thompson, F.L., Thompson, C.C., Swings, J., 2003. Vibrio tasmaniensis sp. nov., isolated from Atlantic Salmon (Salmo salar L.). Syst. Appl. Microbiol. 26, 65–69. 10.1078/072320203322337326

Tourneroche, A., Lami, R., Hubas, C., Blanchet, E., Vallet, M., Escoubeyrou, K., Paris, A., Prado, S., 2019. Bacterial–Fungal Interactions in the Kelp Endomicrobiota Drive Autoinducer-2 Quorum Sensing. Front. Microbiol. 10, 1693. 10.3389/fmicb.2019.01693

Vanhove, A.S., Rubio, T.P., Nguyen, A.N., Lemire, A., Roche, D., Nicod, J., Vergnes, A., Poirier, A.C., Disconzi, E., Bachere, E., Le Roux, F., Jacq, A., Charriere, G.M., Destoumieux-Garzon, D., 2016. Copper homeostasis at the host vibrio interface: lessons from intracellular vibrio transcriptomics. Environ. Microbiol. 18, 875–888. 10.1111/1462-2920.13083

Wang, X., Li, Z., Su, J., Tian, Y., Ning, X., Hong, H., Zheng, T., 2010. Lysis of a red-tide causing alga, Alexandrium tamarense, caused by bacteria from its phycosphere. Biol. Control 52, 123–130. 10.1016/j.biocontrol.2009.10.004

Wang, Y., Li, S., Liu, G., Li, X., Yang, Q., Xu, Y., Hu, Z., Chen, C.-Y., Chang, J.-S., 2020. Continuous production of algicidal compounds against Akashiwo sanguinea via a Vibrio sp. co-culture. Bioresour. Technol. 295, 122246. 10.1016/j.biortech.2019.122246

Weerasinghe, A.J., Amin, S.A., Barker, R.A., Othman, T., Romano, A.N., Parker Siburt, C.J., Tisnado, J., Lambert, L.A., Huxford, T., Carrano, C.J., Crumbliss, A.L., 2013. Borate as a Synergistic Anion for Marinobacter algicola Ferric Binding Protein, FbpA: A Role for Boron in Iron Transport in Marine Life. J. Am. Chem. Soc. 135, 14504–14507. 10.1021/ja406609s

Wu, J., Luther, G., 1994. Size-Fractionated Iron Concentrations in the Water Column of the Western North-Atlantic Ocean. Limnol. Oceanogr. 39, 1119–1129. 10.4319/lo.1994.39.5.1119

Xu, Q., Wang, P., Huangleng, J., Su, H., Chen, P., Chen, X., Zhao, H., Kang, Z., Tang, J., Jiang, G., Li, Z., Zou, S., Dong, K., Huang, Y., Li, N., 2022. Co-occurrence of chromophytic phytoplankton and the Vibrio community during Phaeocystis globosa blooms in the Beibu Gulf. Sci. Total Environ. 805, 150303. 10.1016/j.scitotenv.2021.150303

Yang, Q., Feng, Q., Zhang, B., Gao, J., Sheng, Z., Xue, Q., Zhang, X., 2021. Marinobacter alexandrii sp. nov., a novel yellow-pigmented and algae growth-promoting bacterium isolated from marine phycosphere microbiota. Antonie Van Leeuwenhoek Int. J. Gen. Mol. Microbiol. 114, 709–718. 10.1007/s10482-021-01551-5

Zhang, W., Wang, Y., Lu, H., Liu, Q., Wang, C., Hu, W., Zhao, K., 2020. Dynamics of Solitary Predation by Myxococcus xanthus on Escherichia coli Observed at the Single-Cell Level. Appl. Environ. Microbiol. 86, e02286–19. 10.1128/AEM.02286-19

